# Extensive recombination suppression and chromosome-wide differentiation of a segregation distorter in Drosophila

**DOI:** 10.1101/504126

**Authors:** Zachary L. Fuller, Spencer A. Koury, Christopher J. Leonard, Randee E. Young, Kobe Ikegami, Jonathan Westlake, Stephen Richards, Stephen W. Schaeffer, Nitin Phadnis

## Abstract

Segregation distorters violate Mendelian Inheritance by over-representing themselves in the progeny of carrier individuals and are commonly associated with chromosomal inversions. When distorting alleles are found on sex chromosomes, the progeny of carrier individuals will exhibit skewed sex ratios, as exemplified by the array of *Sex-Ratio* (*SR*) distorting chromosomes found in Drosophila. Because of the strong selective pressures such chromosomes are thought to inflict on genomes, segregation distorters and their associated inversions are expected to experience rapid turn-over. However, the *SR* X-chromosome of *Drosophila pseudoobscura* is found at high frequencies in natural populations, forms stable latitudinal clines, appears to be unsuppressed, and shows evidence of being quite long-lived. Despite being a historically significant and well-studied segregation distortion system, the mechanisms allowing for the long-term persistence of the *D*. *pseudoobscura SR* chromosome remain unclear. Here, we perform a comparative genomic analysis between *SR* and uninverted standard X-chromosomes in *D*. *pseudoobscura* to study its evolutionary history and dynamics. We find a substantial level of differentiation between the *SR* and standard chromosome and estimate that the associated inversions have likely existed for the entire lifetime of the species (*>* 2 million generations). Through direct recombination experiments and population genetic analyses, we infer that this high level of differentiation is maintained by a combination of suppressed recombination and epistatic selection. Finally, our data reveal a massive mutational target size for protein and expression level changes specific to *SR* generated by its three non-overlapping inversions. Together our results imply that the entire *SR* chromosome in *D*. *pseudoobscura* behaves as a single co-adapted gene complex and has been maintained through a combination of suppressed recombination and epistatic selection. This finding highlights the dramatic consequences segregation distorters can have in shaping chromosome-wide patterns of recombination, nucleotide variation, and gene expression.

## Introduction

Unlike normal X-chromosomes, *Sex-Ratio* chromosomes are selfish variants that distort Mendelian segregation ratios in their own favor by destroying Y-bearing sperm. As a result, males that carry a *Sex-Ratio* chromosome produce nearly all female progeny. Such *Sex-Ratio* chromosomes have been found in many Dipteran species [1–4], and are almost always associated with chromosomal inversions. The *Drosophila pseudoobscura Sex-Ratio* (*SR*) chromosome represents one of the longest studied and enigmatic *Sex-Ratio* chromosomes [5–8]. This *SR* chromosome distorts sex chromosome segregation ratios nearly completely, and is found at surprisingly high frequencies in local populations, sometimes at more than 30% frequency, and forms latitudinal clines [5, 6, 9]. When Y-chromosomes are under such constant attack by X-linked segregation distorters, this leads to a powerful selective force for Y-chromosome variants that are resistant to distortion or autosomal alleles that suppress distortion [4, 10, 11]. Despite the strength of distortion and the high frequency that the *D*. *pseudoobscura SR* chromosome can reach within local populations, no resistant Y chromosomes or suppressor alleles have been identified after extensive searches [12].

The *Sex-Ratio* chromosome of *D*. *pseudoobscura*, like most other *Sex-Ratio* chromosomes, is associated with chromosomal inversions [5, 6]. The *D*. *pseudoobscura SR* chromosome carries three non-overlapping inversions on the right arm of the X-chromosome (*XR*) (where the distorting alleles are located) with respect to the wild type or Standard (*ST)* X-chromosome. Segregation distorters are thought to be closely associated with chromosomal inversions because inversions can create groups of tightly linked alleles that evolve in relative isolation from uninverted regions [13–16]. Even the simplest models of distortion require an association between at least two interacting alleles: a driving locus that causes distortion and a responder locus on which the driving locus can act [17, 18]. An association of segregation distorter systems with chromosomal inversions can prove advantageous by preventing recombination between the distorting locus and responder alleles and, thus, avoid the formation of suicide chromosomes *i*.*e*., when the distorter and sensitive responder alleles are found on the same chromosome, leading to self-destruction; [13, 17]. Sex chromosomes, however, do not generally recombine with each other in Drosophila. The prevention of suicide chromosomes may thus explain the association of inversions with autosomal distorters, but is insufficient to explain the association of sex chromosome segregation distorters with chromosomal inversions. Instead, chromosomal inversions may permit sex chromosome segregation distorters to persist by allowing the accumulation of alleles that either enhance distortion or help evade suppressors-of-distortion. According to this idea, distorter systems that become associated with inversions enjoy an advantage by generating stronger drive mechanisms or evading suppression, and hence rise in frequency within populations. Distorting chromosomes that are associated with chromosomal inversions may, thus, evolve as large co-adapted gene complexes that drive efficiently.

Here, we perform a comparative analysis of the *Sex-Ratio* (*SR*) and *Standard* (*ST)* strains of *D*. *pseudoobscura* to uncover the evolutionary history of the distorting chromosome. We first mapped and sequenced the precise breakpoints for two of three of the *SR* chromosomal inversions, which indicate that the direct physical position effects of the inversions are unlikely to underlie the *SR* phenotype. Second, we estimated the divergence of the *SR* chromosome using sequences that flank the inversion breakpoints, and find that it likely arose around the time that *D*. *pseudoobscura* itself originated, suggesting that these inversions have persisted for the entire lifetime of the species. Third, we examine the population genetics of *SR* by analyzing single nucleotide polymorphisms (SNPs) across the chromosome, and find that the chromosome is highly differentiated and distinct from the *ST* chromosome. Surprisingly, this pattern of differentiation spans all three non-overlapping inversions as well as a large section that is collinear between the *SR* and *ST* chromosomes. Fourth, through direct experimentation we show that recombination can and does occur in this collinear region, albeit at severely reduced rates, indicating that recombination suppression extends well beyond inversion breakpoints and epistatic selection is required to maintain high differentiation. Finally, we show that this high differentiation has led to a large number of fixed amino acid changes and a significant enrichment of differentially expressed genes across the right arm of the X-chromosome. Our results imply that the entire *SR* chromosome in *D*. *pseudoobscura* behaves as a single co-adapted gene complex and has been maintained through a combination of suppressed recombination and epistatic selection.

## Results

### Identification of chromosomal inversion break-points

To investigate the population genetics of the *Sex-Ratio* chromosome in *D*. *pseudoobscura*, we collected wild flies from Zion National Park, UT and screened them for males that display strong sex ratio distortion. We isolated stably-distorting stocks that produce *>*95% female progeny, and confirmed the presence of the three *SR* associated inversions with polytene chromosome analyses (Figure 1). The *D*. *pseudoobscura* chromosome contains three non-overlapping inversions on the right arm of the X chromosome (*XR*, Muller element A D): basal, medial, and terminal. Previously, the breakpoints of these inversions were coarsely mapped to major sections on the polytene maps (basal: section 23D to 24D; medial: section 25D to 34A; and terminal: section 39A to 42B) [6, 19]. We pooled DNA from eight independent *SR* lines and eight matched *ST* lines, Illumina sequenced them, and realigned the paired-end reads to the *D*. *pseudoobscura* reference genome (*v*.*3*.*2*). We used the coarse locations on the chromosomal maps to precisely determine the coordinates of the inversion breakpoints on the physical map [20]. In particular, we searched for Illumina read-pairs from the *SR* strains that aligned in the same orientation, yet in different regions of the chromosome separated by large distances (*>*1Mb) [21, 22]. By scanning through these aberrantly mapped reads, we were able to identify candidate positions of two of the three pairs of inversion breakpoints.

**Figure 1:**
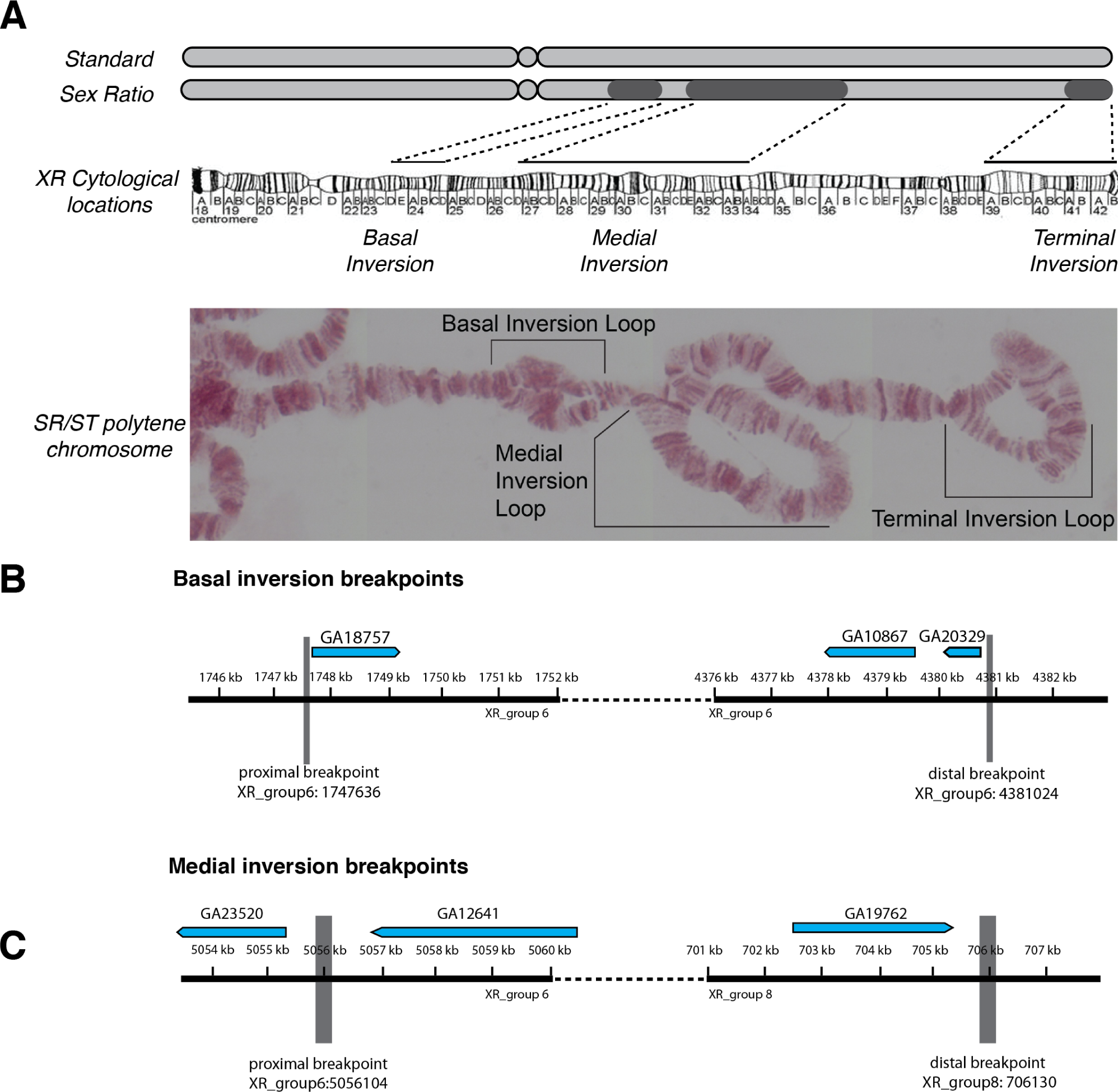
The structure of the *Sex-Ratio* (*SR*) chromosome and inversion breakpoints. **(A)** A schematic of the standard (*ST)* and *SR* X-chromosomes, with the darker regions showing the approximate locations of the basal, medial, and terminal inversions. The dotted lines show the locations of the three nonoverlapping inversions on the cytological diagram and polytene image of a *SR*/*ST* heterozyote female. **(B)** The genomic location and size of the basal inversion breakpoints. The coding regions of adjacent genes are shown above. **(C)** The genomic location and size of the medial inversion breakpoints. Similarly, the coding regions of adjacent genes are shown above.

To confirm the identity of the inversion breakpoints, we designed PCR primers to amplify the breakpoint regions only when the region is inverted relative to a *ST* chromosome (*i*.*e*., the reference sequence). Using this approach, we were able to confirm both the proximal and distal breakpoints of the basal and medial inversions. By amplifying the breakpoints and Sanger sequencing the amplicons, we were able to locate and genetically characterize precise molecular breakpoints of the basal and medial inversions (see Supplementary Material for locations and sequences). We were, however, unable to precisely locate either of the break-points of the terminal inversion due to its proximity to the telomere, which consists of large blocks of repetitive sequences. Therefore, we use approximate cytological locations for the terminal inversion breakpoints in all subsequent analyses.

The basal inversion breakpoints are nearly precise, and contain only a small insertion of genetic material at the proximal breakpoint. The proximal breakpoint of the medial inversion is similarly precise, but interestingly, when we used BLAST to identify the location of the breakpoint sequence, it aligned not only to this breakpoint, but also to many other locations in the genome. Transposable element insertions have been previously implicated in the formation of inversions through homologous exchange of sites on either side of the inversion [23–25]. The presence of a potential repetitive element at these inversion breakpoints may indicate that this inversion formed through similar processes. Moreover, the distal breakpoint of the medial inversion not only contained this same sequence, but nearly 10 kb of sequence not found in the reference genome. These results show that the sequences at the breakpoints of the basal and medial inversions are quite different from each other, possibly reflecting different mechanisms of their formation.

### **Estimating the age of the *SR* chromosome and its three non-overlapping inversions**

Because recombination is most restricted between arrangements in regions near the inversion breakpoints, these regions preserve their evolutionary history in their patterns of genetic divergence [22, 26–31]. We used the regions flanking the breakpoints (±250 kb) to estimate the divergence between the *SR* and *ST* arrangements to determine their age (see Supplemental Information Tables S1-2 for details on the next generation sequence alignment statistics). Across these regions, we estimated *F*_*ST*_ and absolute sequence divergence (*d*_*xy*_) in 10 kb non-overlapping windows [32–34]. Between the *SR* and *ST* arrangements we observed high overall levels of differentiation with a mean *F*_*ST*_ of 0.225 (95% CI: 0.211-0.241). When we compared the emphD. pseudoobscura *ST* chromosome with the outgroup *D*. *miranda*, we observed a higher mean *F*_*ST*_ of 0.403 (95% CI: 0.388-0.418) in these same regions. Using the classical transformation of Cavalli-Sforza [35] and scaling to a speciation time of 2 million-year-ago (Mya) between *D*. *pseudoobscura* and *D*. *miranda*, we estimate this corresponds to a divergence time of 0.99 Mya (95% CI: 0.92-1.07) for the *SR* and *ST* arrangements. This falls within the upper range of the divergence time estimate obtained by Babcock & Anderson [8] and precedes the divergence of *D*. *pseudoobscura* and its sister species *D*. *persimilis* [8, 31, 36]. Between *SR* and *ST* in these regions, the mean *d*_*xy*_ was 6.55 × 10^−3^ (95% CI: 6.17-6.93 × 10^−3^) while between *D*. *miranda* and *ST, d_xy_* was estimated as 1.03×10^−2^ (95% CI: 0.99-1.06×10^−2^), indicating that divergence is significantly greater (Mann-Whitney *U* test, *p <* 2 × 10^−16^) between the two species than between the two arrangements. The *D*. *pseudoobscura SR* chromosome, thus, appears to have originated after the split from *D*. *miranda*, but before or close to the divergence time between *D*. *pseudoobscura* and *D*. *persimilis* [31].

We next compared estimates of *F*_*ST*_ and *d*_*xy*_ for each inversion individually to infer the order of their formation on the *SR* chromosome. *F*_*ST*_ does not significantly differ (Mann-Whitney *U* test, *p <* 0.519) between the regions surrounding the basal (*F*_*ST*_ : 0.262, 95% CI: 0.231-0.292) and medial (*F*_*ST*_ : 0.251, 95% CI: 0.232-0.271) inversion breakpoints. Likewise, *d*_*xy*_ does not significantly differ (Mann-Whitney *U* test, *p <* 0.281) between the regions surrounding the basal (*d*_*xy*_: 7.01×10^−3^, 95% CI: 6.24-7.98×10^−3^) and medial (*d*_*xy*_: 6.68×10^−3^, 95% CI: 6.14-7.32×10^−3^) inversion breakpoints. However, in regions flanking the terminal inversion breakpoints, both *F*_*ST*_ (mean: 0.162, 95% CI: 0.143-0.183) and *d*_*xy*_ (mean: 5.67×10^−3^, 95% CI: 5.31-6.13×10^−3^) are significantly lower (Mann-Whitney *U* test, *p <* 2×10^−16^) compared to the medial and basal breakpoints, providing evidence that the terminal inversion is the youngest on the *SR* chromosome. Although the terminal inversion appears to be younger than either the basal or medial inversions, the overall high levels of divergence and differentiation suggest it is still quite old. Using the same transformation of *F*_*ST*_, we estimate the age of the terminal inversion to be approximately 662 Kya (95% CI: 578-754), which is at the lower end of the range of the divergence time estimated by Babcock & Anderson [8] and still predates the divergence of *D*. *pseudoobscura* and *D*. *persimilis* [8, 36]. Rather than the *SR* chromosome gradually accumulating inversions sequentially over its lifetime, our results point towards the rapid formation of the *SR* chromosome, with all three associated inversions present in the ancestral species or soon after the split with *D*. *persimilis*, similar to what is observed for the single *SR* associated inversion in *D*. *persimilis* [31].

### **Extensive genetic differentiation across the three non-overlapping *SR* inversions**

We extended our analysis of differentiation by estimating *F*_*ST*_ in 10 kb non-overlapping sliding windows across the entire length of *XR* to test if the high differentiation observed flanking the inversion break-points was unique to these regions. Previous experimental and theoretical studies have suggested that differentiation decreases towards the center of large inversions as the result of double crossover recombination events within inversion heterozygotes [26, 37]. For the basal and medial inversions, we observe a significant (*p*_*Basal*_ *<* 0.013; *p*_*Medial*_ *<* 2 × 10^−16^), albeit weak 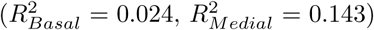, negative linear relationship between the distance to the nearest inversion breakpoint and *F*_*ST*_. Interestingly, *F*_*ST*_ actually significantly increases (*p*_*Distal*_ *<* 1.02×10^−14^) with distance from the breakpoints for the distal inversion, although this relationship is still fairly weak 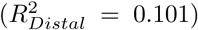. Overall, differentiation does not dramatically decrease within the center of the inversions, as the average *F*_*ST*_ is 0.247 (95% CI: 0.233-0.259) and 0.120 (95% CI: 0.105-0.137) for the central 500 kb of the basal and medial inversions, respectively. Together, this suggests that although some low level of exchange may occur through gene conversion or double crossovers, high levels of genetic differentiation are not restricted to the breakpoints and instead remain elevated across each inversion.

The three non-overlapping inversions of the *SR* chromosome are separated by large intervening collinear regions (totaling 5.6 Mb of the chromosome), where single or double crossovers can potentially form and lead to a reduction in the levels of genetic differentiation relative to inverted segments. Instead, we observe a chromosome-wide effect of extensive genetic differentiation, with *F*_*ST*_ elevated for a majority of the chromosome arm across these collinear regions (Figure 2). Even in the large collinear regions separating the inversions, *F*_*ST*_ is not significantly less than in regions contained within the three *XR* inversions (Mann-Whitney *U* test, *p <* 0.99; Figure 3).

**Figure 2:**
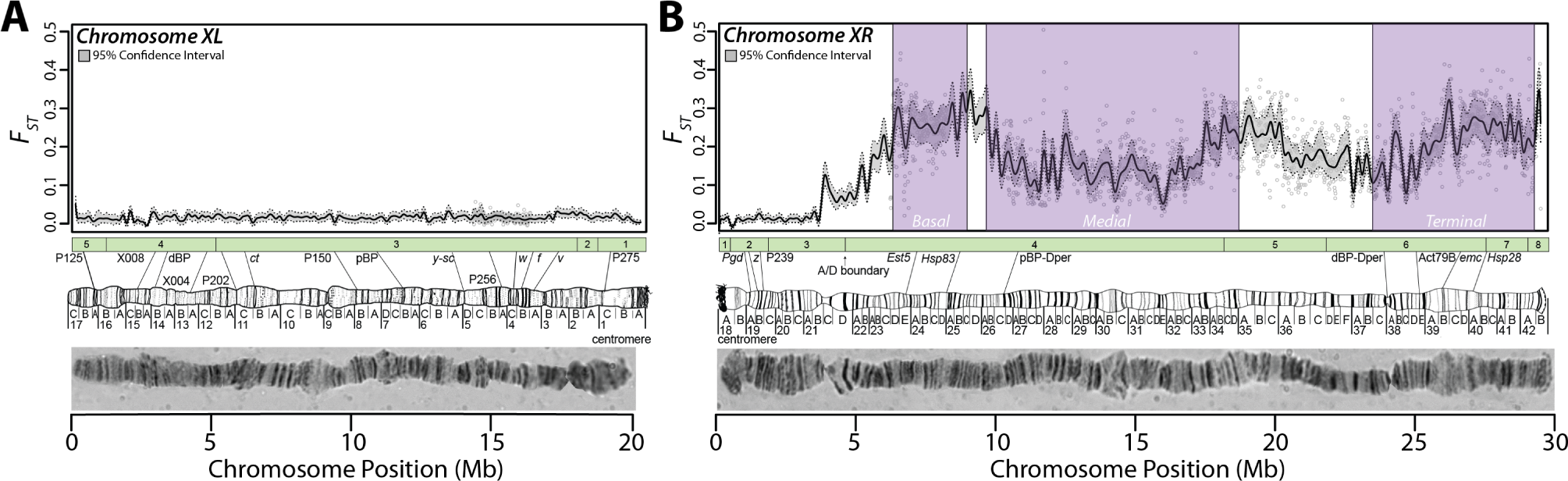
Elevated genetic differentiation across *XR*. *F*_*ST*_ was estimated in 10 kb non-overlapping sliding windows across chromosomes *XL* **(A)** and *XR* **(B)**. Gray dots show the estimate of *F*_*ST*_ in each window. The black line represents the loess smoothed average *F*_*ST*_ across the chromosome and the gray region bounded by dotted lines is the loess smoothed average 95% bootstrapped confidence interval of the mean estimate of *F*_*ST*_ within each window (note this differs from the 95% confidence interval estimated across regions discussed in the main results). On *XR*, purple shaded regions indicate the locations of the basal, medial, and terminal inversions. Polytene images of each chromosome and sketches of the cytogenetic regions with the approximate locations of common genetic markers are depicted below each plot. The green boxes represents the linear ordering and size of genomic scaffolds used to construct the chromosome sequence from Schaeffer *et al*. [20].

**Figure 3:**
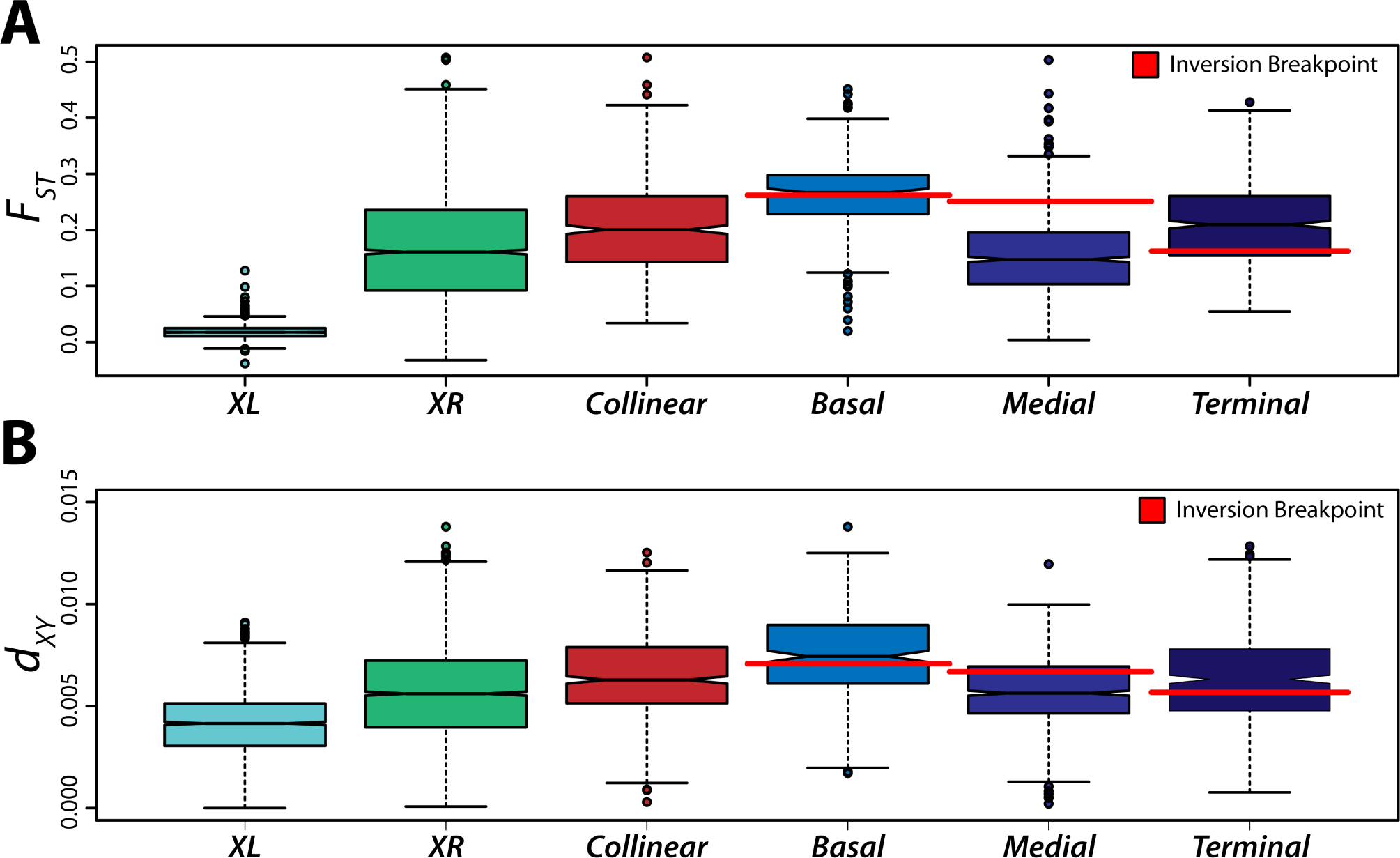
The distribution of differentiation and divergence estimated across genomic regions. Differentiation (*F*_*ST*_, **A**) and divergence (*d*_*xy*_, **B**) were estimated in 10 kb non-overlapping sliding windows across *XL* and *XR* and within the three inverted regions and intervening collinear regions on *XR*. The boxplots depict the distribution of each statistic estimated for each genomic region. The red lines indicate the mean of each statistic estimated for regions flanking (*±* 250 kb) the inversion breakpoints for that boxplot.

### Genetic differentiation is caused by recombination suppression and epistatic selection

To determine if the increased genetic differentiation observed across *XR* is primarily driven by the presence of inversions on the chromosome and not by residual population structure or X chromosome effects, we then estimated *F*_*ST*_ across *XL*, which does not harbor any inversions and thus recombination occurs freely. We observed substantially lower levels of differentiation on *XL* (mean *F*_*ST*_ : 0.018, 95% CI: 0.017-0.018; Figure 3). Moreover, significantly lower differentiation is also observed in the proximal region of *XR* approaching the basal *SR* inversion (mean *F*_*ST*_ : 0.055, 95% CI: 0.049-0.0610) compared to the other collinear regions of the chromosome arm (Mann-Whitney *U* test, *p <* 2×10^−16^), where recombination is also free to occur. Hence, it appears unlikely that population structure is a major influence of the increased *F*_*ST*_ and *d*_*xy*_ observed within inverted and intervening collinear regions on *XR*, as it appears gene flow is unimpeded in other regions of the X chromosome. Instead, we hy-pothesized this extensive genetic differentiation across the chromosome could be due to either completely suppressed recombination in these regions or epistatic selection acting on linked inversions.

A cytogenetic analysis of 107 and 96 offspring from two female heterozygotes (ST/SR) found no evidence for recombination between the medial and terminal inversions (see Supplemental Information for methods and results). These data suggested that if crossing over happens between the medial and terminal inversions, it occurs at a frequency *<* 1% (Supplemental Table S3-4), but a more sensitive assay was needed to detect lower recombination frequencies. Thus, we performed additional recombination experiments using three independently sampled *SR* chromosomes isolated on a multiply marked standard arrangement genetic background. The isogenic stock for background replacement carried mutants of *sepia* (*se*^1^, 156.5 m.u. marking the basal and medial inversions) and *short* (*sh*^1^, 225.9 m.u. marking the terminal inversion); therefore recombination, or lack thereof, can be directly assayed with standard testcrossing procedures (Figure 4). The collinear region of 4,745,273 bp between the medial and terminal inversions is *>* 50 cM on the standard genetic map [38], and models of genetic flux with inversions incorporating interference suggest crossover rate in this region should be 0.01-0.001 events per meiosis [26, 39]. Consistent with these interference models, rare recombinants have sometimes been observed in nature [19, 40].

**Figure 4:**
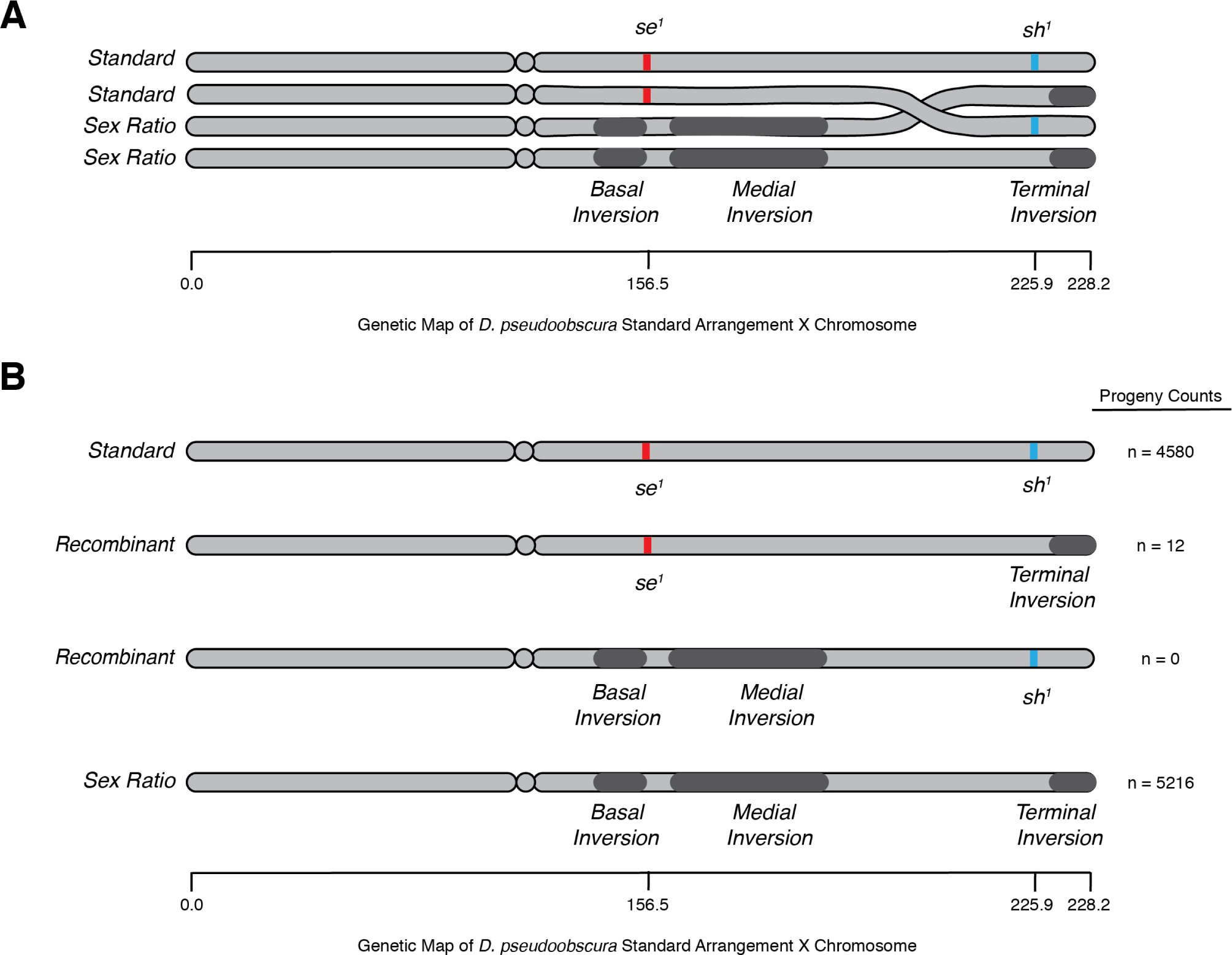
Overview of recombination experiments. **(A)** Illustration of the four strand bundle present in prophase of Meiosis I for a *SR*/*ST* heterozygote. Depicted in red and blue are the relative position of visible mutations to inversions of the *SR* chromosome (∼70 cM on the standard arrangement genetic map). The position of both markers and inversions relative to the standard arrangement genetic map is approximate and not exact because inversion heterozygosity strongly distort this map. **(B)** The four possible chromosomes recovered in the recombination experiment, with the pooled progeny counts recorded to the right, please note the complete absence of one complementary class of recombinants (basal and medial inversions with visible marker *sh*^1^).

In the recombination experiment a total of 10,891 progeny were scored from 33 experimental bottles, ten replicate bottles for each of three *SR* isolates and three replicates of a single *ST* gene arrangement. The recombination fraction observed between *se* and *sh* in *ST* arrangement homozygotes was 0.4224; with Kosambi’s [41] correction this translates to a genetic distance *>* 50 cM, consistent with previous observation [38]. From all *SR*/*ST* heterozygotes only 12 recombinant chromosomes were recovered, yielding an estimated genetic distance of 0.12 cM, indicating a roughly 500-fold decrease in recombination in the collinear region of SR chromosomes (Figure 4; Supp. Table S5). This recombination fraction is consistent with the lower values predicted by interference model of recombination suppression for inversion heterozygotes [26, 39].

Although recombination in the collinear region is strongly suppressed, it is not completely eliminated. The 0.0012 rate of crossing-over is sufficient, especially over long periods of time, to cause dissociation of the terminal inversion from the basal and medial inversions of *SR* chromosomes, with linkage equilibrium being established in under 10,000 generations. Interestingly, in this experiment 12 of 12 recombinants carried only the terminal inversion, while none of the complementary class (basal and medial inversion only) were recovered, a very unexpected result 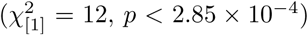 (Supp. Table S6). Furthermore, none of the terminal inversion carrying recombinants were observed to distort sex ratio. This suggests that in addition to strong recombination suppression, there is also epistatic selection acting on the linked inversions to maintain a co-adapted gene complex resulting in the elevated genetic differentiation across all inversions and intervening collinear regions of *XR*.

### **The *SR* inversions behave as a single evolutionary unit and generate extensive LD across *XR***

The non-overlapping *SR* inversions effectively suppress recombination across a large majority of the *XR* chromosome, behaving as a single massive rearrangement rather than three independent inversions. Here, we predict suppressed recombination to have generated a strong signature of LD and long-range associations across *XR*, both within inverted segments and in the intervening collinear regions. Although we can analyze allele frequencies across the genome from our pooled libraries, we cannot assemble individual haplotypes. Thus, to examine linkage disequilibrium between segregating sites across the chromosome, we designed PCR primers to amplify eight intergenic regions on *XL* and *XR* (Supplemental Table S6). We sequenced these regions from all eight strains of both *SR* and *ST*, and concatenated the sequences to perform the LD analysis (Figure 5). Of all valid pairwise comparisons between segregating sites, 10% of them show significant LD [42]. Furthermore, within each intergenic region on *XR*, there is at least one site that shows a significant association of alleles with the *SR* arrangement (Fisher’s exact test, *p <* 0.05), yet no site on *XL* shows a similar association. Moreover, each of the *XR* regions distal to the basal inversion contains at least one site that is in significant LD with a site from each of the other more distal regions. This signature of LD is consistent with the pattern we observed using *F*_*ST*_ in which the *SR*-specific differentiation is found not only within the inversions, but also in the intervening collinear regions, further supporting the suppression of recombination across the majority of *XR*.

**Figure 5:**
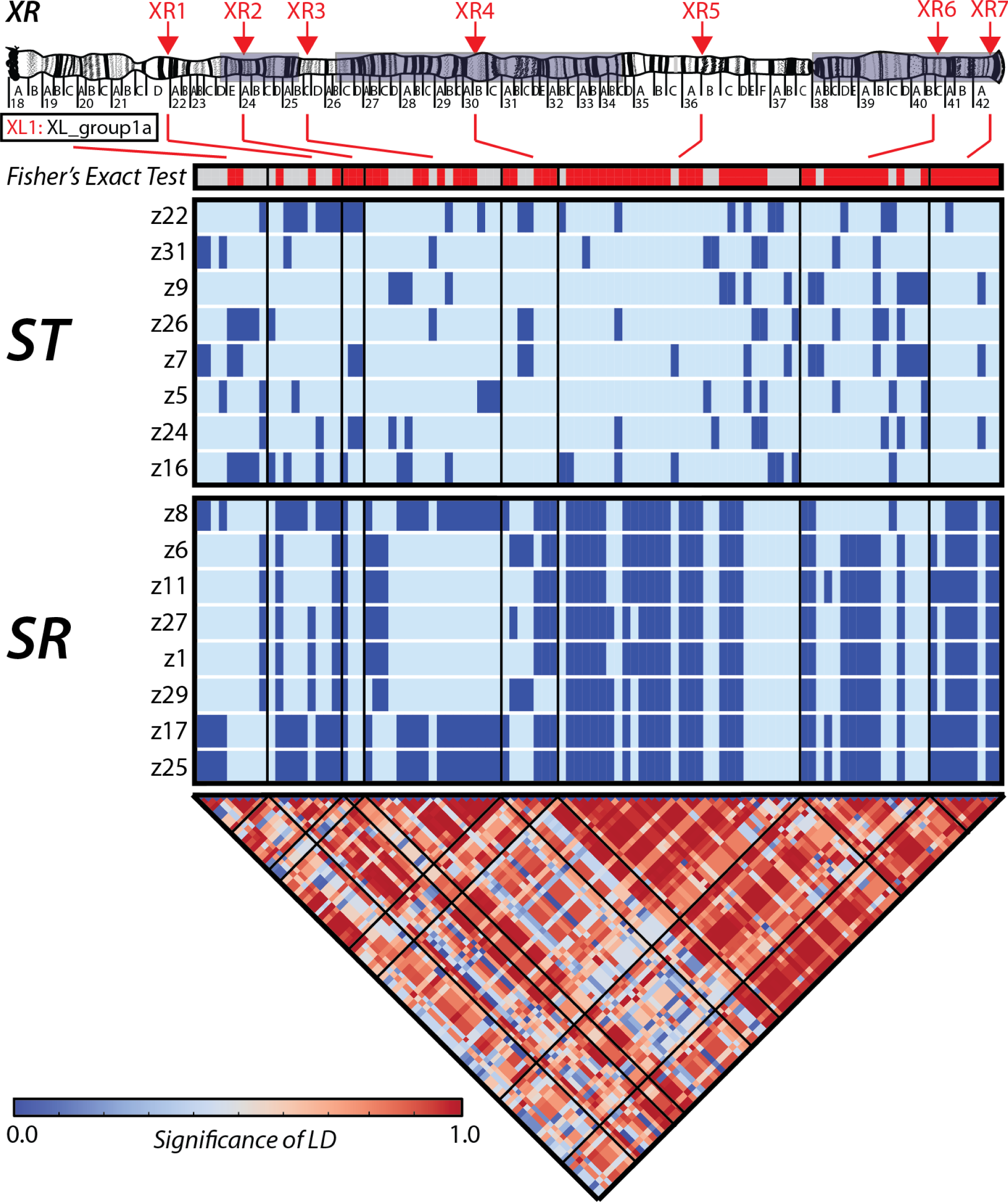
Long-range linkage disequilbirum is present across XR. Linkage disequilibrium (LD) was estimated using the PCR amplified sequences of 8 intergenic markers. The red arrows at the top of the chromosome sketch show the approximate location of each sequenced marker. The single horizontal bar depicts the results (*p <*0.05=red, *p <≥*0.05 gray) of a Fisher’s exact test for the association between alleles and chromosome (*SR* and *ST)* type. The following horizontal bars show haplotype diagrams for polymorphic sites in the sequenced *ST* (top) and *SR* (bottom) strains, with darker colored sites representing the derived allele. The bottom triangular heat map shows the significance of LD for all polymorphic sites in the sequenced intergenic regions estimated with the correlation-based approach of Zaykin *et al*. [43]. Red indicates greater LD and blue represents non-significant allele associations. The black lines show the boundaries between each intergenic region.

The analysis of sequences from PCR amplicons allows us to test for the significance of LD between intergenic regions. Alternatively, by analyzing those sites that are fixed in our pooled sample of *SR* chromosomes, yet not observed in *ST*, we can examine the pattern of alleles that are in perfect LD with one another. After determining the ancestral state with *D*. *miranda*, there are a total of 45,849 sites which are fixed for a derived allele unique to *SR* across *XR* observed in our sample. We next estimated the proportion of such fixed derived sites in 10 kb windows, finding an astonishingly high proportion with a chromosome wide mean of 1.55×10^−3^ (95% CI: 1.49-1.62×10^−3^). In other words, on average, for a given 10 kb interval more than 15 sites are fixed within our sample of *SR* chromosomes for a derived allele that is not observed in *ST*. This proportion is even higher within inverted regions of the chromosome, with a mean value of 1.80 × 10^−3^ (95% CI: 1.72-1.89 × 10^−3^). Furthermore, the proportion of such sites does not significantly differ between inverted regions and the collinear segments that separate them (Mann-Whitney *U* test, *p <* 0.066). In the 10 kb window with the largest proportion of these sites, found within the basal inversion, more than 1% of all base pairs are fixed for an *SR*-specific derived allele. These results indicate the *SR* arrangement behaves as a single evolutionary unit, with long-range associations of alleles and large numbers of derived sites held in complete linkage disequilibrium.

### **The *SR* inversions create a massive mutational target size for protein and expression level changes**

The largest fraction of differentiated and derived sites on *SR* chromosome are likely simply due to approximately one million years of evolution without gene flow, however a subset of these sites held together in perfect association may lead to functional changes and are potential candidates for the *SR* phenotype. Furthermore, it is also possible that a few of the subset of functional changes may also be responsible for mitigating the effect of suppressors, allowing the *SR* trait to persist for longer than would be expected under a model of rapid suppression. By taking advantage of the massive recombination restricted environment of the *SR* inversions, these suppressors-of-suppressors may have been able to evolve alongside the distorting alleles allowing them to evade any genomic mechanisms that may arise to limit them. To identify potential targets for both the evolution of distorters and their enhancers, we first sought to determine the genes which contain fixed amino acid differences. Of the total number of derived sites fixed in *SR*, 8,156 occur within protein coding regions, including 2,773 nonsynonymous changes found across 983 genes. This corresponds to over 35% of all genes on *XR* containing at least one fixed amino acid difference between *SR* and *ST*. In contrast, there are only 71 derived sites that are fixed in *SR* and not observed in *ST* in total across *XL*, none of which fall within a protein coding gene. Meanwhile, 549 genes on *XR* contain multiple fixed amino acid differences. Interestingly, the gene (*GA28653)* with the largest number of fixed amino acid changes (21) is the ortholog to *Spc105R* in *D*. *melanogaster*, which produces a kinetochore protein that is required for the co-orientation of sister centromeres during meiosis and promotes the accurate segregation of chromosomes [44]. These results indicate that multiple genes may be involved in the *SR* phenotype, yet also suggest large amounts of potentially functional amino acid changes unrelated to segregation distortion may be tightly linked to the causal mutations as a consequence of suppressed recombination and high divergence of the chromosome.

A total of 13 fixed sites specific to SR are predicted to be loss-of-function mutations because they introduce premature stop codons or disrupt splice sites, including in the *D*. *melanogaster* orthologs of *Gale, RAF2*, and *CG17744*. Interestingly, in *D*. *melanogaster* complete loss of *Gale* expression is lethal and the gene plays a key role in the metabolism of dietary galactose [45]. In our *SR* sample, a premature stop codon is introduced in the third exon that is predicted to have a high functional impact. Although further experimental work would be needed to functionally describe and estimate the fitness effects of these predicted loss-of-function and amino acid altering mutations, these results indicate that a substantial number of potentially protein altering changes are harbored on the SR chromosome, some of which may even have deleterious effects on fitness.

In addition to amino acid changes, functional effects may result from changes in patterns of gene expression. We, therefore, performed RNA-seq to test for significant expression level differences between *ST* and *SR* (see Supplemental Tables S7-8 for RNA-Seq read data and differential expression statistics). Genome-wide, a total of 1,203 genes were identified as significantly (*q <* 0.05, BH corrected) differentially expressed. A total of 868 of these genes are located off of the X chromosome. Although these differences may be the result of *trans-* acting factors associated with genetic variation located on the X chromosome, our crossing scheme maintained lines through different sets of marker strains, possibly creating structured variation on the autosomes. Additionally, the *SR* and *ST* strains carry different third chromosome arrangements (Arrowhead for *SR* and Standard for *ST)* which are known to harbor expression differences [46]. Indeed, a principal component analysis (PCA) of SNPs called in autosomal genes from the RNA-seq reads confirmed the presence of variation distinguishing *ST* and *SR* individuals (Supplementary Figure S3). It is therefore possible that these autosomal transcriptional differences may arise from *cis-*acting factors associated with structured genetic variation in the stock marker stains and as a result, we restrict our analysis to the 335 genes detected as differentially expressed on the X chromosome (Figure 6A). Significant differences in expression were detected for 43 genes on *XL* and for 292 genes on *XR*. The proportion of differentially expressed genes on *XR* is significantly greater than on *XL* (Fisher’s Exact Test: *p <* 1×10^−5^). For differentially expressed genes on *XR*, there is an enrichment of those upregulated (177) relative to *ST* than those that show lower expression (115; Fisher’s Exact Test: *p <* 0.009). Inverted regions contain 198 differentially expressed genes, while the collinear regions separating them contain 84. However, the proportion of genes that are differentially expressed within the inversions (.122) is not significantly different than the proportion of differentially expressed genes found in the collinear regions (.133; Fisher’s Exact Test: *p <* 0.510), consistent with the effects of suppressed recombination and differentiation extending beyond the breakpoints and across the chromosome arm.

**Figure 6:**
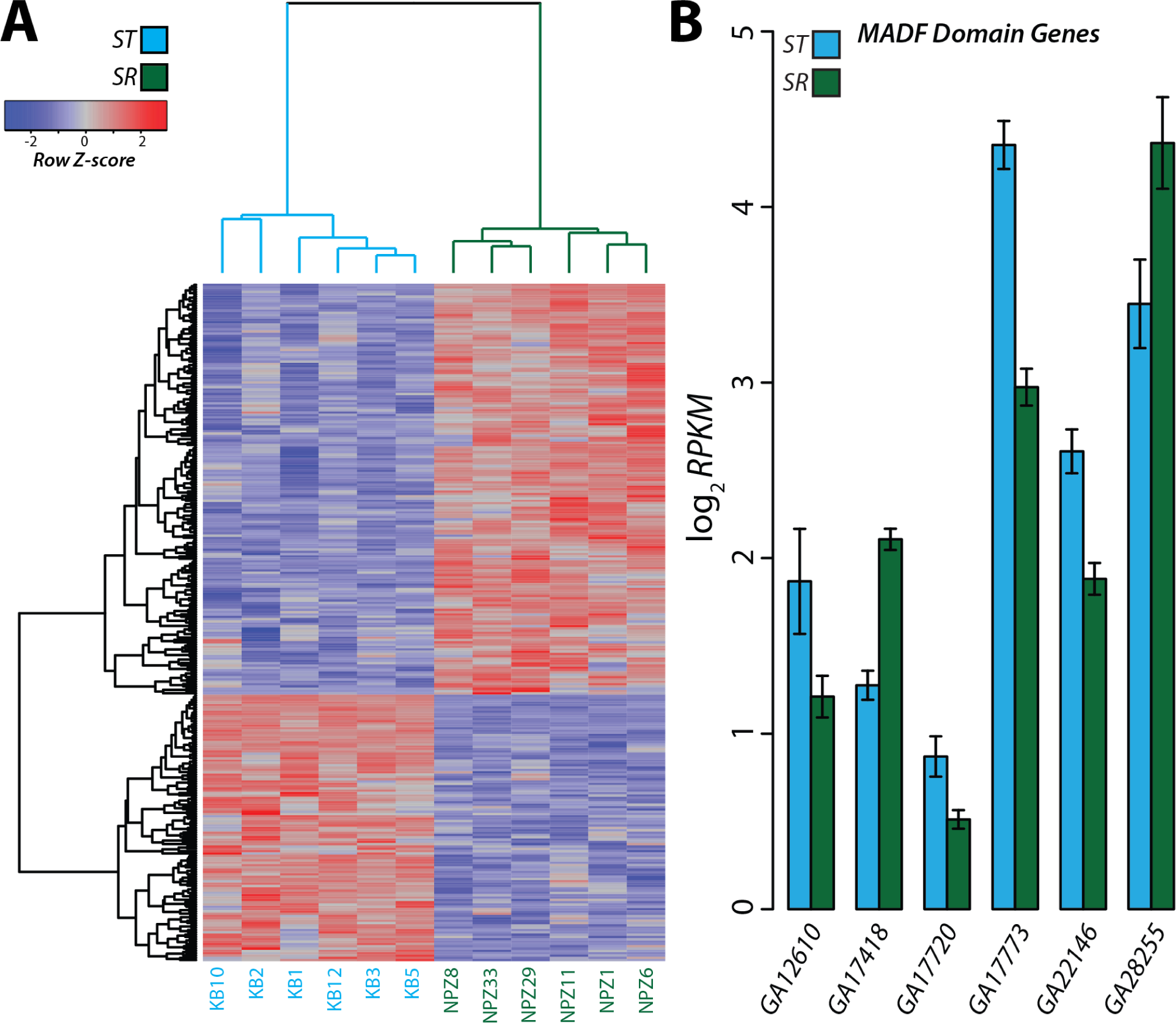
Differentially expressed genes across *XR*.] **(A)** Differential expression depicted as a heat map for the 292 significant (*q <*0.05) genes on *XR*. The individuals strains (columns) and genes (rows) are arranged according to unsupervised hierarchical clustering. Each gene is colored according to the deviation from the mean level of expression across all individuals. **(B)** Expression levels for the 6 MADF domain containing genes detected as significantly differentially expressed. The height of each bar represents the mean expression for *ST* and *SR* measured as reads per kilobase per million mapped reads (RPKM) with error bars indicating the standard error.

We performed a gene ontology (GO) analysis using DAVID software to test for the enrichment of common biological functions, pathways and protein domains among genes we detected as differentially expressed [47]. After correcting for multiple testing, the only category that remains significantly enriched among the 335 differentially expressed genes on the X-chromosome contain MADF domains (*q <* 0.041, BH corrected). This category contains 6 genes (Figure 6B) that are all located on *XR*. In fact, the significance increases if only the 292 differentially expressed genes located on *XR* are used as input (*q <* 0.032, BH corrected). Four of the genes (*GA17720, GA17773, GA22146, GA28255)* also harbor multiple fixed amino acid changes unique to *SR* with the *D*. *melanogaster* orthologs to *CG11723* and *stwl* containing 15 and 17, respectively. Intriguingly, MADF domain containing proteins are also implicated in hybrid-incompatibilities in *D*. *melanogaster* and its closely related species *D*. *mauritiana, D*. *simulans*, and *D*. *sechellia* [48–51]. No-btably, another gene product containing a MADF domain is *Overdrive* (*Ovd)*, which has previously been identified as a single gene that underlies both male sterility and segregation distortion in hybrids between the USA and Bogota subspecies of *D*. *pseudoobscura* [52]. Here, although we did not observe differential expression in *Ovd*, our results further implicate MADF domain containing proteins in the phenomenon of segregation distortion as we see a significant enrichment for this group of genes among our set of differentially expressed transcripts. In combination with the previously described role of MADF domains in segregation distortion, this set of differentially expressed genes are attractive candidates for follow-up studies to dissect the molecular basis of the *SR* trait.

## Discussion

The *Sex-Ratio* system of *D*. *pseudoobscura* has served as an important example of segregation distortion and its impact on the evolution of species. Despite long-standing hypotheses that unsuppressed systems are a fleeting stage of the lifetime of a distorting system, our results support that the *SR* chromosome is in fact quite old [7, 8, 53]. Previous phylogenetic analyses of the *Esterase-5* region have indicated an ancient monophyletic origin of the *SR* chromosome, suggesting it was segregating in the ancestral species of *D*. *pseudoobscura* and *D*. *persimilis* and diverged from *ST* at least 2 million generations ago [8]. Other inversions elsewhere in the genome, such as several polymorphic third chromosome arrangements, are similarly ancient in *D*. *pseudoobscura* demonstrating that old segregating inversions are a common feature of this species [8, 22, 30]. Segregation distorters and their associated inversions are predicted to rapidly turn over within populations rather than exist at stable equilibria over long evolutionary timescales because of the extreme selective pressures they exert on populations [11, 54, 55]. Here, we used polymorphisms surrounding the recombination restricted areas around inversion breakpoints to estimate their ages and found that all three inversions arose at least 660 Kya, in agreement with previous estimates [8]. Our results show that the inversions associated with segregation distorters are so old they may, in fact, predate the divergence of the species that carry them and is similar to recent studies which highlight similar ancient origins of other selfish meiotic drive elements, such as the *t*-haplotype in mice [56, 57].

Through direct estimation of recombination rates and high-resolution analysis of genetic variation, our results support a combination of recombination suppression and epistatic selection acting to maintain the *SR* chromosome inversions as a co-adapted complex. Considering the equilibrium frequency of *SR* chromosomes observed in natural populations of D. pseudoobscura (*q ≤* 0.30) and strongly suppressed recombination in the collinear regions (*r* ≈ 0.001), given these parameters and in the absence of selection, the recombinant chromosomes observed in our experiments should reach a frequency of *q*(1–*q*) at linkage equilibrium. The neutral expectation for decay of linkage disequilibrium (*D*) between the medial and terminal inversions of *SR* chromosomes can be described with the recursion equation *D*_*t*_ = *D*_0_(1–*r*)*^t^*, indicating dissociation of the terminal inversion should be complete within 5,000 generations (*t*). A more explicit deterministic model that assumes equilibrium frequencies of *SR* chromosomes are a result of drive-selection balance, as well as incorporating the sex-limited recombination of Drosophila sex chromosomes, shows the approach to linkage equilibrium is somewhat slower but still achieved within 10,000 generations, which is still substantially less than the 2,000,000+ generations since the formation of the *SR* chromosome. Furthermore, the deterministic model indicates that recombinant *SR* chromosomes, or at least the terminal inversion only, should be sampled up to 20% in present day natural populations. In contrast to these simple expectations, only five recombinant *SR* chromosomes have ever been sampled in the 80+ years of natural population surveys and we detect no evidence of recombination in the *F*_*ST*_, *d*_*xy*_, or LD data. The expected behavior of *SR* chromosomes, even with 500-fold suppressed recombination, in conjunction with observations of natural population, genomic analysis of differentiation, and an estimated age of at least 660 thousand years indicates the presence of a strong evolutionary force, most likely epistatic selection, acting the *SR* chromosomes. We hypothesize strong suppression of recombination generated by the three *SR* inversions across the majority of the chromosome may create favorable conditions for epistatic selection to arise and allow for the persistence of the chromosomes in populations longer than expected under models of rapid turnover or neutrality.

Here, our results support the extensive increased differentiation is a result of a combination of suppressed recombination and epistatic selection across the three non-overlapping inversions. Increased differentiation and divergence extending beyond inversion breakpoints into collinear regions has previously been observed on the third chromosome of *D*. *pseudoobscura* and several other Drosophila species [22, 36, 58–60]. Several previous studies have demonstrated that this effect is amplified in systems with non-overlapping inversions and noted similar drastic reductions from the expected frequencies of recombinants [61–64]. Consistent with observations in *D*. *subobscura* and *D*. *ananasae*, our results show that crossovers are strongly suppressed between non-overlapping inversions even when they are separated by large blocks of collinear regions [61, 63].

The *D*. *pseudoobscura SR* system is one of several that defies the logic on the age of distorting systems within Drosophila [31, 65, 66] and stands in contrast with theory proposing that distorting systems should be short-lived [11, 54]. As no suppressors of *D*. *pseudoobscura SR* have been identified, it is unclear how the trait is maintained as a stable polymorphism over the long evolution of this species. It has previously been proposed that additional secondary loci may arise within the inversion containing the primary distorting alleles that counteract the effect of suppressors found elsewhere in the genome [3, 67]. Here, we show that by strongly suppressing recombination across the intervening collinear regions, the three *SR* chromosome inversions are transmitted as a single large inversion, thereby creating a massive mutational target in which enhancers of drive or “suppressors-of-suppressors” could arise. When this strong reduction in recombination is coupled with epistatic selection to maintain inversion linkage, the long-term evolutionary persistence of the *SR* chromosomes has allowed for the formation of a highly-differentiated region spanning across three non-overlapping inversions. Direct experimentation demonstrates that the recombination suppression effects of inversions extend well beyond their immediate breakpoints and this can be further compounded in the case of multiple, sequential inversions. Here, the three inversions have generated a single evolutionary unit spanning more than 80% of *XR* and containing more than 2,100 genes. Held together in this massive region of extensive divergence are a large number of functional changes, including more than 500 genes with multiple derived amino acid differences in perfect LD and more than 200 significantly differentially expressed transcripts. While the genes underlying the *SR* phenotype are almost certainly contained among these, our results may also contain candidate loci which have become associated with the primary distorting alleles because they impede the action of suppressors or act as enhancers.

By completely distorting against the competing Y chromosome, the *D*. *pseudoobscura SR* chromosome ensures its transmission into the progeny of carrier males. In the absence of suppressors, this allele would be expected to rapidly sweep to fixation, eliminating males and extinguishing carrier populations. However, the advanced age of this chromosome, coupled with its relatively stable population frequencies during the last century of sampling and apparent lack of suppressors, indicate other evolutionary forces must act to prevent it from sweeping to fixation. It has been argued stable population frequencies could be the result of competing evolutionary pressures: reduced reproductive fitness in carrier males may balance the advantage gained through selfish distortion by the *SR* chromosome [68, 69]. For example, fewer offspring are produced by *SR* males carriers in populations of *D*. *simulans*, although similar findings in *D*. *pseudoobscura* are less conclusive [70, 71] (see Supplemental Table S4). However, *D*. *pseudoobscura* female re-mating experiments suggest that *SR* males have reduced sperm competitive ability relative to *ST* males and thus the prevalence of *SR* in nature may be maintained by polyandry [68, 69, 72].

While it could also be tempting to attribute the evolutionary persistence of *SR* to a single closely-linked deleterious allele that strongly reduces its fitness, any chromosome uncoupled from a deleterious allele would rapidly increase in frequency. However, if multiple deleterious alleles become linked with the distorter, they could prevent its disassociation and reduce its overall fitness. In the case of *D*. *pseudoobscura SR*, the massive differentiated region of *XR* is home to thousands of genes, a large number of which contain at least one fixed missense mutation or are differentially expressed, some of which may slightly lower fitness and together balance the meiotic advantage of the *SR* chromosome. Thus, the large mutational target created by the *SR* inversions that may harbor the counter-suppressive alleles, may also create a large opportunity for the accumulation of deleterious mutations. It has long been known that segregation distortion is capable of shaping the genetic content of inversions that contain them, and can allow for the persistence of alleles that would otherwise be purged by selection. In fact, we identified multiple loss-of-function variants and potentially deleterious missense mutations fixed within the *SR* background, for example in the ortholog of *Gale*. Future studies aimed at measuring the fitness effects of *SR* in natural populations will be critical in distinguishing between the evolutionary forces which contribute to the maintenance of segregation distortion in *D*. *pseudoobscura*.

There are at least three possible explanations for the existence of unsuppressed *Sex-Ratio* chromosomes. First, a *Sex-Ratio* chromosome may have arisen recently, with not enough time for the evolution of resistant Y chromosomes or autosomal suppressors. Contrary to this idea, previous estimates of the age of the *D*. *pseudoobscura SR* chromosome, although based on sequences from a limited number of loci, suggest the inversions associated with the *SR* trait may be surprisingly old and likely arose at least 2 million generations ago [8, 54, 73, 74]. Second, a sex chromosome distorter may remain unsuppressed if there is insufficient genetic variation within natural populations. *D*. *pseudoobscura*, however, is known to harbor ample genetic variation across the genome and particularly with Y-chromosomes [22, 53, 75], suggesting that the absence of suppressors is unlikely to be limited by natural genetic variation. Third, *Sex-Ratio* chromosomes and suppressors may be locked in an ongoing evolutionary arms race, with the distorting chromosome rapidly evolving enhancers of distortion or alleles that allow the evasion of suppressors-of-distortion. Thus, the current unsuppressed state may only be a brief transitory phase in the long-term evolution of *SR* chromosomes. Although intuitive, there is so far little genetic evidence to support or reject this scenario.

Our results show that the three nonoverlapping inversions suppress recombination across a vast majority of the chromosome arm and combined with the action of epistatic selection, effectively operate as a single, large inversion by limiting exchange even in the intervening collinear regions separating their breakpoints. By suppressing recombination, the *SR* inversions have generated an extensive and massive region of divergence and differentiation, with loci held in significant LD across megabase scales. This extensive level of differentiation has created a large mutational target for potential enhancers of distortion, suppressors-of-suppressors, or linked deleterious alleles to arise and evolve alongside the *SR* segregation distorter. Thus, despite being one of the longest studied selfish chromosomes, many fundamental genetic and evolutionary aspects of the *D*. *pseudoobscura SR* chromosome has remained mysterious [5]. Our results now provide insight into the evolutionary mechanisms acting to maintain the *SR* chromosome and that have allowed it to persist for a substantially long period of time.

## Materials and Methods

### **Collection, isolation and maintenance of *Sex-Ratio* chromosome strains:**

We collected wild *D*. *pseudoobscura* flies from Zion National Park, UT, USA in September 2013 using bait consisting of an assortment of rotten fruits and screened them for the presence of *Sex-Ratio* chromosomes. Individual wild males collected were crossed to females from a *Standard D*. *pseudoobscura* stock with multiple markers on the X-chromosome: *cut*^1^ (*ct*^1^, 122.5), *scalloped*^1^ (*sd*^1^, 1 43), *yellow* (*y*^1^, 174.5) and *sepia*^1^ (*se*^1^, 1145.1) [38]. Males carrying a *Sex-Ratio* chromosome are readily identified as those that produce nearly all female progeny. To screen for *Sex-Ratio* chromosomes in females, we allowed individual wild-caught females to produce progeny in the laboratory. The resulting sons were individually crossed to *ct, sd, y, se* females. Males carrying *Sex-Ratio* chromosomes were similarly identified as those that produced nearly all female progeny. We bred and and tested a total of 113 *D*. *pseudoobscura* individuals, consisting of 66 males and 47 females. Of the 66 males collected and screen, 5 of these males had an *SR* chromosome. Of the 47 females collected, 10 carried an *SR* chromosome. Of 160 *D*. *pseudoobscura* X chromosomes tested (66 from males, 94 from females), 145 were *ST* chromosomes and 15 were *SR* chromosomes; *i*.*e*., *SR* chromosomes we found at a frequency of approximately 9.4% in this population. Once *SR* males were identified, we generated homozygous *SR* females using the sepia marker, which is known to cover the basal inversion on the *SR* chromosome [8]. All stocks were raised on standard cornmeal media at 18° C.

### DNA extractions and sequencing

To generate whole genome shotgun sequencing libraries for *D*. *pseudoobscura* strains, we first pooled one male each from 8 *SR* strains and 8 *ST* strains from our Zion National Park collections. We extracted DNA from these flies using the 5 Prime Archive Pure DNA extraction kit according to the manufacturers protocol (ThermoFisher, Waltham, MA). All libraries were generated with the Illumina TruSeq Nano kit (Epicentre, Illumina Inc, CA) using the manufacturers protocol, and sequenced as 500bp paired end reads on a Illumina HiSeq 2000 instrument.

### Sequence alignment and SNP identification

Low-quality bases were removed from the ends of the raw paired end reads contained in FASTQ files using seqtk (https://github.com/lh3/seqtk) with an error threshold of 0.05. Illumina adapter sequences and polyA tails were trimmed from the reads using Trimmomatic (*v0*.*30)* [76]. The read quality was then manually inspected using FastQC. Following initial pre-processing and quality control, the reads from each pool were aligned to the *D*. *pseudoobscura* reference genome (*v3*.*2)* using bwa *v0*.*7*.*8* with default parameters [77]. Of the total reads, 95.82% and 94.87% mapped successfully for the *ST* and *SR* pools, respectively (Supplementary Table S1). Genome wide, the average fold coverage was ∼74× and ∼75× for the *D*. *pseudoobscura ST* and *SR* pools, respectively. For X-chromosome scaffolds, the average fold coverage was ∼45× and ∼46× (Supplementary Table S2).

After the binary alignments were sorted and indexed with SAMtools (*v0*.*1*.*19)* [78], single nucleotide polymorphisms (SNPs) were called using freebayes (*v0*.*9*.*21)* [79] with the expected pairwise nucleotide diversity parameter set to 0.01, based on a previous genome-wide estimate from *D*. *pseudoobscura* [80]. The samples were modeled as discrete genotypes across pools by using the “–J” option and the ploidy was set separately for X chromosome scaffolds (1*N)* and autosomes (2*N)*. SNPs with a genotype quality score less than 30 were filtered from the dataset. Across the genome we identified a total of 3,598,524 polymorphic sites, 751,556 and 634,610 of which were located on chromosomes *XR* and *XL*, respectively. Sequences are deposited on the NCBI Short Read Archive (SRA) with accession numbers SRR6331544 & SRR6331545.

### Identifying and confirming the inversion break-points

We located the inversion breakpoints for first two inversions of the *D*. *pseudoobscura SR* chromosome by viewing the mapped paired end reads of the *ST* and *SR* pooled genome sequences in the Integrated Genomics Viewer application using two methods. 1) We interpret the mapped paired end reads by pair orientation, such that parallel mapped paired end reads where the read pair is mapped farther than expected and in the same orientation in the *SR* sequence but not the *ST* sequence is a clear indication that an inversion breakpoint is present. 2) Our sequencing library was prepared using 500 bp paired end reads. When mapped paired end reads are located approximately 500 bp from each other in the *ST* strains, but map over 1 Mb in *SR* strains this is a clear indication that an inversion breakpoint is at that location.

Inversion breakpoints were confirmed molecularly through a polymerase chain reaction (PCR) inversion assay. For proximal breakpoints, the forward primer is common to *ST* and *SR* with the reverse primer unique to *ST* or *SR*. For distal breakpoints, the forward primer is unique to *ST* or *SR* and the reverse primer is common to both *ST* and *SR*. For primers unique to *SR*, they were designed approximately 500 bp from the opposite inversion breakpoint (if designing for proximal breakpoint, primer designed 500 bp before distal break-point).

### Estimates of differentiation and divergence

To estimate population differentiation we used the Reich-Patterson estimator of *F*_*ST*_ [34], which has previously been shown to be unbiased and have higher accuracy than other estimators when the sample size is small but the number of markers is large [81]. Within each pool, we first calculated the frequency of the alternative allele at each site. *F*_*ST*_ was then directly estimated from the resulting allele frequencies. Sites that contained missing data were not considered. We estimated *F*_*ST*_ in 10 kb windows and obtained 95% confidence intervals for the chromosome region of interest by performing 10,000 bootstrap replicates across each. A negative *F*_*ST*_ value indicates greater differentiation within a population than between populations. Divergence time estimates were taken with the Cavalli-Sforza [35] transformation of *F*_*ST*_ as

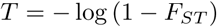

and then multiplied by a scaling factor in each window so that the divergence time between *ST* and *D*. *miranda* was 2 Mya [31].

Absolute sequence divergence was estimated with *d*_*xy*_, a measure of the mean number of pair-wise nucleotide substitutions [32, 33]. 95% confidence intervals were similarly obtained by performing 10,000 bootstrap replicates across each region of interest. Custom Python code used to estimate *F*_*ST*_ and *d*_*xy*_ as well as all R scripts used for plotting and statistical analyses are available at https://github.com/zfuller5280/Dpse_SR_analyses.

### **Analysis of linkage disequilibrium of the *D*. *pseudoobscura SR* chromosome:**

As a result of our pooled sequencing design, individual haplotypes could not be constructed from the assembled Illumina reads. Therefore, we designed PCR primers (Supplemental Table S3) to amplify intergenic regions located on *XL* and inside and outside of the inversions on *XR*. The chromosomal locations and approximate coordinates of the sequences are:

**XL1** XL;

XL group1a:2,958,187-2,959,179.

**XR1** proximal of basal;

XR group6:370,850-371,767.

**XR2** inside basal;

XR group6:3,450,538-3,451,504

**XR3** distal of basal/proximal of medial;

XR group6:4,760,237-4,761,215

**XR4** inside medial;

XR group6:9,392,822-9,393,842

**XR5** distal of medial/proximal of distal;

XR group8:2,908,477-2,909,427

**XR6** inside terminal;

XR group3a:327,359-328,353

**XR7** distal of terminal;

XR group5:349,989-350,987

We amplified the intergenic regions of 8 *ST* strains and 8 *SR* strains using the polymerase chain reaction. We then directly Sanger sequenced the amplicons using the same primers. The sequences for each of the regions were aligned, and indels and singletons were removed for the analysis of linkage disequilibrium (LD). Segregating sites from each region were concatenated into a single sequence and LD was estimated using the correlation based method of Zaykin *et al*. [43]. For each site, we also performed a Fisher’s exact test to determine the significance of allele association with *ST* or *SR*. Significance values were corrected for multiple testing using the Benjamini-Hochberg procedure [82].

### Analysis of recombination rates between medial and terminal inversions

To directly test for recombination in the collinear region between medial and terminal inversions of *D*. *pseudoobscura SR* chromosomes we conducted a series of well-controlled testcrosses. Three independent *SR* chromosomes sampled from Zion National Park were isolated and background replaced by a minimum of 7 generations of backcrossing to an isogenic stock. This isogenic stock carries the visible mutations sepia1 (*se*^1^, 1-145.1 marking the basal and medial inversions) as well as *short*^1^ (*sh*^1^, 1-225.9 marking the terminal inversion)[38], and has undergone *>* 7 generations of full-sib mating in the Phadnis Lab prior to use in experimental crosses.

The recombination experiments follow the standard mapping conditions of Bridges [83] modified for the life-history and reproductive biology of *D*. *pseudoobscura*. In this case, 20 virgin females heterozygous the markers were collected over a seven day period, aged for an additional seven days, crossed to 20 males of the tester strain (*se*^1^ *sh*^1^) under light CO_2_ anesthesia, allowed 24 hours to recover, and then tap transferred into milk bottles with 50 mL of standard cornmeal-molasses medium. Egg laying period lasted seven days, after which adults were removed from bottles and 0.5% propionic acid was used to hydrate food as necessary. Emerging progeny were scored for visible markers daily starting from day 20 until the last individuals eclosed, only male progeny were scored because variable expression of the wing vein mutation *sh*^1^ was observed in females. The experiment was conducted at room temperature without controlling for relative humidity or light/dark cycle.

The recombination experiment was conducted as a single block, fully randomized design, with experimenter blind to treatment. A total of 33 experimental bottles were setup, consisting of ten replicate bottles for each of the *SR* chromosome isolates in the heterozygous state and three additional bottles with *ST* /*ST* heterozygotes to calibrate our estimated genetic distances under these experimental conditions. The recombination rates for *se* and *sh* in the standard arrangement are so high, that after correcting for interference and multiple crossover events with Kosambi’s function [41], the genetic map distance exceeds the maximum limit of detection in a two point testcross (*>* 50 cM). In contrast, the extremely low recombination rate from all ten bottles for each *SR* chromosome isolate required the data was pooled and reported with an exact binomial 95% confidence interval. All recombinants, as determined by visible markers, were subsequently confirmed by scoring the presence/absence of the medial and terminal inversions of *SR* chromosomes via polytene chromosomes squash, and a chisquare test for complementary classes of recombinants was conducted using the 1:1 Mendelian expectation.

### RNA Collection

We isolated RNA from testes of 6 biological replicates of *SR* and *ST* fly strains. For each biological replicate we pooled tissue dissected from between 40-50 individuals. Individuals for each strain were maintained in three separate technical replicate growth chambers containing standard cornmeal-agar-molasses food media with yeast. The pooled tissue was immediately snap-frozen in liquid nitrogen after dissection and stored at −80°C prior to RNA extraction. RNA was purified with RNeasy spin-columns (Qiagen) using the manufacturers instructions and stored at −80°C before performing RNA sequencing. Total RNA concentrations for each sample were quantified using a nanodrop (Thermo Scientific).

### RNA-Seq

Illumina RNA-Seq (Wang et al. 2009) was performed following standard protocols by the Baylor College of Medicine Human Genome Sequencing Center, Houston, TX on an Illumina HiSeq 2000 sequencing platform. Briefly, poly-A+ mRNA was extracted from 1 *µ*g total RNA using Oligo(dT)25 Dynabeads (Life Technologies, Cat. No. 61002) followed by fragmentation of the mRNA by heat at 94°C for 3 minutes (for samples with RIN=3 - 6) or 4 minutes (for samples with RIN of 6.0 and above). First strand cDNA was synthesized using the Superscript III reverse transcriptase (Life Technologies, Cat. No. 18080-044) and purified using Agencourt RNAClean XP beads (Beckman Coulter, Cat. No. A63987). During second strand cDNA synthesis, dNTP mix containing dUTP was used to introduce strand-specificity. For Illumina paired-end library construction, the resultant cDNA was processed through end-repair and A-tailing, ligated with Illumina PE adapters, and then digested with 10 units of Uracil-DNA Glycosylase (NEB, Cat. No. M0280L). Amplification of the libraries was performed for 13 PCR cycles using the Phusion High-Fidelity PCR Master Mix (NEB, Cat. No. M0531L); 6-bp molecular barcodes were also incorporated during this PCR amplification. These libraries were then purified with Agencourt AMPure XP beads after each enzymatic reaction, and after quantification using the Agilent Bioanalyzer 2100 DNA Chip 7500 (Cat. No. 5067-1506), libraries were pooled in equimolar amounts for sequencing. Sequencing was performed on Illumina HiSeq2000s generating 100-bp paired-end reads. RNA-Seq Accession Numbers in the SRA database: (ST Biosample Numbers: SAMN06208344-SAMN06208349; SR Biosample Numbers: SAMN06208350-SAMN06208355).

### Read Mapping and Analysis of Differential Gene Expression

The reads generated from RNA-Seq were mapped to the D. pseudoobscura reference genome (*v3*.*2)* using the subjunc aligner (*v1*.*4*.*6)* under default parameters [84]. As recommended in the users manual, read ends were not trimmed before aligning to the reference genome because the software soft clips ends with low mapping quality (MAPQ) scores. In total, over 755 million read pairs were generated. Between 33.2 million and 96.8 million reads were produced for each individual replicate (Supplemental Table S4). An average of 81.9% of reads mapped to annotated features in the *D*. *pseudoobscura* reference genome. There was not a significant difference in the fractions of reads that mapped successfully between *SR* or *ST* replicates (82.0% and 81.7% respectively). Using featureCounts (emphv1.4.6), the number of reads mapping to each annotated exon were counted. We filtered out genes that did not have a minimum of 10 reads mapped in at least three individuals. After removing genes from the data that did not meet our filtering criteria, 14,687 genes were retained for analysis. 2,247 of these genes are located on scaffolds mapping to *XR*, while 2,298 are on scaffolds mapping to *XL*.

After filtering, upper-quartile between-lane normalization was performed using the R package EDASeq [85]. The read counts were further normalized using the *RUVs* method implemented in RUVSeq [86]*RUVs* is a normalization procedure to control for unwanted variation not associated with the biological covariates of interest (here, *SR* or *ST)* in the data. The factors of unwanted variation were estimated from the genes within each replicate group (*ST* and *SR*) because no differential expression is expected. Normalization factors were estimated using the “relative log expression” (RLE) method of [87].

Differential gene expression was investigated in the normalized read counts using the R package edgeR (*v3*.*10*.*2)* [88]. The covariates of interest (*i*.*e*., X chromosome arrangement) and the first factor of unwanted variation (*k*=1) were used to construct the design matrix of the negative binomial generalized linear model (GLM). Briefly, the GLM takes the form of

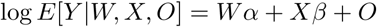

where *Y* is the matrix containing the read counts for each gene, *W* is the matrix containing the factors of “unwanted variation”, *X* is the matrix containing the covariates of interest and *O* is a matrix of offsets estimated through upper-quartile normalization. *α* and *β* indicate the parameters for the factors of unwanted variation and covariates of interest (*i*.*e*., “treatment effect”, here the X chromosome arrangement) respectively.

To test for significant differential expression between *ST* and *SR* males, a quasi-likelihood (QL) F-test was performed as implemented in edgeR with the *glmQLFTest()* function. The QL F-test is preferred to a standard likelihood ratio test because it reflects the uncertainty in dispersion estimates for each gene and is a more robust and reliable method to control for the error rate [89]. To correct for multiple testing, we corrected the raw *p*-values using the Benjamini-Hochberg method [82]. We considered genes with a false discovery rate (FDR) *<* 0.05 as significantly differentially expressed (see Supplementary Table S8 for a complete list of raw and corrected *p*-values for all genes).

