## Supporting Information for "Extensive recombination suppression and chromosome-wide differentiation of a segregation distorter in Drosophila"

**Supplementary text: Sequences at the breakpoints of the basal and medial inversions**

**1. Basal inversion, proximal breakpoint**
XR_group6:1747336-1747636-BREAKPOINT-XR_group6:4381020-4380840

GCACGTCACMCAATACAAACACACAAGCAGTTATTCGACTAACAAATCACTTTCTTTTCACATTTCACTTGT
GCTTGGCTGTACATTCGATTACATGGGTATTTTGTTGCTCAAAACTTGTATTTATTTCTTTATATAGTATACA
TTACAGTGGTTCAAGTGCATATTTTTTTTGTGTGCGAATATTTTATTTGCAAAAAAGTAGAGATGGAGAAAC
GTCGATTTTACTATCGATAAATACTCAGCCGGCTGGTAGCGATGTCTATCGATCCTACGCTGCCAGATCGA
GAGAAAACGGTTACCTATATCTTAACCATCTGTATGGTTAGAAACAAGACAAGACAAGGCAAAACTTTCCC
ATCTCTAAGAAAGTATAATATGATATTTACAAATTCTTTGAAAAAGATAACTTCAGTGCGTACCAAACACTAA
TTGACTTGATTGACGAAATTGATGTAAAACTATAACCGGTACTCCAACACCTTAAAAAAAGCGGGCGGTCA
AGTAATCCCACGCTACTCAATGGTAGCATGGTAACTTAAGCCCTC

**2. Basal inversion, distal breakpoint**
XR_group6:1747856-1747736-BREAKPOINT-XR_group6: 4381024-4381264

ATAGGGATGTTAAGTATTCATTGATATCAATTATATGTATATGTTTTGCATATCATCATCGATATTTTTGTGC
GATAAATATTATTTCGATTGAAATTGTAAATACCTAATGATGTATTTCAATTATTTTCGCGGTATATTCTGCT
GATTTTTTGTATTTTAAGGCACTAGCACCACTAACCATATCCAGTGGGATATATACTGTATATATATATATAC
CATACAGTATACTGTATGGTTAGTGGTGGTACAATGGCGCTATCGATACTATCGCATGCATCACCTATGTT
ACTAGGAACTACATACATTTTTTCGAGTGTGTGCCTATCAAATGCGTTACCGTCTGTTGATTCTTTAATCTT
TAAGTTTAGTTTTATTTGGGAAGCACGGTTTGTACGATGTTGAGCGCAAAAGCGAGAGAAAACCTCCATCG
AATGACTCTCCCGCGTTATGTAATTAAACGCATTGTTCAGTTGCTCCTCACTCAACTGG

Repetitive motif at breakpoint: CCATACAGTATACTGTATGGTTAGTGG

**3. Medial inversion, proximal breakpoint**
XR_group6: 5055855-5056104-BREAKPOINT-XR_group8: 706130-705579

ATTACCGGAGAATATTTAAATTTAATACGTTCCAGCGATAAATATACCGATTCGAGTAATAGCGAGTTGGAT
AAAATAATAACACAATTAGAAATGTGTATATCTATGTGTTTTTTTATTTTAAGCCTATTTTTTGGCCTTTTTCA
ATTTTTAGCTTGTTTTAGCTTGTGTTTTTGTGAAATCTAGCTTATTTTTACGCTCAAAATCTGGCAACACTGC
TGCAGTAGTGACCGCGGATTTATCGATAACGACCATCAGGAATATACTGAAACATACCATCTCATTTTAAA
AATATACCGTAAATATACTGACGAATTCAAGTTCTATTTTACTTATTTCTCGTTTTTGATATTTCGTCGAATAT
TACCAGCTATATAGAACATTTAGCCATGCCCACTCAATTTTGTACGATTGATGAATCAATTCTATACATGAT
TGGCCTATTCGAAATACTTGCTTTTATTGGATTTTGGCTAAAACAAGGCATAAACAAAATAGTTCAACAAAA
AGAAACAGTGTTAAGATAGCAAGGTGTATTAATATAAGGTAACGTGTACAAGGTGAACAGTGTGTGCACTC
CAATGGTTATCCTTTTTTAGAGGATTATCGATTTTCGATAAAAACCTTTACTGGTCTTTTTCGAACTCAAATC
AAATTTTAATATATAAATTGCTAGTTAACTTTAAATTGTTGAACTTTATTTCTGTTAGCTTACAATATACATAA
TTCAAAATGTTTTAGGCCCACATACGAATGAAATGTGGGAAAATTGGTTTTACATGAATAGAATTCACCATT
TTATAAATTCTTACAATTTTCGTGTTTTCTTTTTTGTTTTTTCTTCACTTAACCTTAACTGGAACCGAGAGCG
AAAGTAGAAAAACATTCAATTTCTTAAACAGTAGCGATTTTTTGTTGTAGTTAAATATTCGTGTATGTCCCGC
CTAGGAATTCTTCCGATATTATAACTTTTTGAGAGAGATTTAGATCAAATAGTAGACATACAGATACATATA
CATATACATATATTTCATATATGTATGTGTATATA

Repetitive motif at breakpoint:
TCAGAAATATACCGAAATATACCATCTCATTTAAAAAATATACCGTAAATATACTGACGAATTCAAGTTCTCT
TTTACATATTCCTCGTTTTTGATCTTCCGTGGAATATTCTCAGCTATATAGAACATTTAGCCATGCCCACTC
AATTTTGTACGATTGATGAATCAATTCTATACATGATTGGCCTATTCGAAATACTTGCATTTATTGGATTT

#### 4. Medial inversion, distal breakpoint

XR_group6: 5058390-5057728,5057645-5056460-BREAKPOINT-XR_group8: 706177-708097

GTTGTACTATAATATATATTGGCAGGCCTTTTGGGTTCGGCATTCGATTCGCATACCCCATAACCATAGCC
ATAACCATACTATATCTTATGGCTGCCATTAAGTTTCATTATTTCTGAATTTGTTTATCCGTACAGGACATCA
ATCAGACACGCCTGCCCCTCATTCCCGAGTATGCCGTGCAGACGCTTACGCCGCAGGAAATGCTTCAGG
TGGAGCAGCGAATGGATCAGCGGAGGGAGCGACTGAAGGACAAGTGCTCGGCCTATGGCCTGGATGTG
TTAGGTGGGTCAATGGGCCACAGCCACAGAGAGAGATAGATAAATAAATAGAGAGAGAAAGAGAGGAAG
AGTGAGAGTGTGAAAACGAACTTCTCCTTCACATCAATCTTATTTACAGGTCACGACTCGTGGCACACCCC
AAACACATGGGAGTTTTTGGTCAACAAAAAGTATCACATTATATGGTGCGTATACAAATTACACATCACATC
ACATTCACAGGAACTCCTCCACTAAAGCCACGAATCTTGCGTAGGTGCAATGTGTTTAAGGCTGCCTCCTC
ATCGTGGATGTTCAACTTCAATGTTCTGGCCGGTTACTCACCCAGTTATTTGCGCAAAACCAAAAAGATTC
TCCTGAATCTGGCCAGAGAACGCTATCCGAGAGTGACCCTTGACGAGGTGAGTGCGGAAATGGAACGGA
AACTAGCATCATACATTTTTTATACCCGATACTCAAAATGAGTATTGGGGTATATAAGATTTGTGGTAAAAG
TGGATGTGTGTAACGTCCAGAAGGAATCGTTTCCGACCCCATAAAGTATATATATTCTTGATCAGCATCAA
TAGCCGAGTCGATAGAGCCCTGTCTGTCTGTCCGTCCGTCCCCTTCACCGCCTAGTGCTCAAAGACTATA
AGAGCTAGAGCAACGATGTTTTGGATCCAGACTTCTGTGATATGTCACTGCTACAAAAATATTTCAAAGAG
CAACCAAATTTGGTATCCACACTCCTAATATATCGGACCGAGACGAGTTTGTTTCAAAATTTCGCCACACC
CCCTTCCGCCCCCGCAAAGGATGCAAATCTGGGGATATTCACAAATCTCAGGGACTATTAAGGCTAGAGT
AACCAAATTTGGTATCCGCACTTCTGCTAGATCTCACTATAAAACGTATATCTCAAAATTTCGCCCCACCCC
CTTCCGCCCCCACAAAGGACGAAAACCTGTTGCATCCACAATATTGCACATTCGAGAAAACTAAAAACGCA
GAATCATAGATAATGACCATATCCATCAGATTGCTGAATCTGGATCACATCAGACAATTTTTATAGCCAAAA
GGAACAAATCAATTTGCACTGGCTACGCAGCGCCCGACGTCACGCTCAGACTGATTTTCTGTCTCTCTCG
CACGCACTCTTTGTCGTGTCGTTTAATATTAGTGGCGTCTGCCGGAGGAGAGCCATACTGACTTAGTATC
GGGTATAACTGTAGAGTTGCGGTGTCCGCTCATAACTCATAACGTTCCCCCTCGTTTTATACCCGATACT
CAAAATGAGTATTGGGGTATATTAGATTTGTGGTAAAAGTGGATGTGTCTAACGTCCAGAAGGAATCGTTT
CCGACCCCATAAAGTATATATATTCTTGATCAGCATCAATAGCCGAGTCGATTGAGCCCTGTCTGTCTGTC
CGTCTGTCCGTCCGTCCGTCTGTCCGTCTGTCCGTCCCCTTCAGCGCCTAATGCTCAAAGACTATAGAGC
TAGAGCAACGATGTTTTTGGATCCAGACTTCTGTGATATGTCACTGCTCGAAANATATTTCAAACTTTGCCC
GCCCACTCCGTCCCCCACAAAGGGCGAATCTGTGCATCCACATTTCGACAATACGAGAAAACTAAAAAAC
GCAGAATCGTAGAAGATGACTATATCTTACAGAGTGCAAAATCTGAACCAGATCGTATAATTATTACAGCC
AGAATCACGAAAACAATTTCACTCTTTCTCGCTCTGTCTCACTCTAACACACAGGTTTCATGGTCGGTTTTG
CCAATTTCAAAATATGAGTTCAAGGATCTCAGAACCTATAAAAGCCAGAGCAACCAAATTTGGTATCCACA
CTCCTGTGATATCGGACCTTGACCGTTTCATGTCCACATTTCGCCACATCCCCTTCCGCCCCCGCAAAGG
ACGAAAATCTGAGGCAACCACAAATCTCAGAGACTATTAAGGCTAGAGTAACCAAATTTGGTACACACACT
CCTTTAAGATGTCACTATAAAACGTATATCTCAGAATTTCGCCCCACCCCCATCCGCCCCCACAAAGGACG
AAAATCTGTTGCATCCACAATATTGCAGATTTGAGAAAACTAAAAACGCAGAATCATAGATAATGACCATAT
CTATTAGATTGCTGAATCTGGATCAGATCAGATAATTTTTGTAGCCAAAAGGAACAAATCAATTTGCATTGG
CTACGCAGCGCCCGACGTCACGCTCAGACTGATTTTCTGTCTCTCTCGCACGCACTCTTTGTCGTGTCGT
TTAATATTAGCGGCCTCTGCCGGAGGAGAGCCATACTGACTTAGTATCGGGTATAACCGTAGAGTTGCGG
TGTCCGCAGCAACTCACAACGTTCCCCCTCGTTTTATGTATCTCGTATGACATTTTTGCTTTTCGTTTGCAG
CTGCGCGAGGCGCAGAATGACTCGCTAACCTTTATCATTGCACGCAATCCCTTTGAGAGGCTGTTGAGCG
CTTATCGCGATAAGATGGTGTTCGCCCTGCCCTATTCCTTCCATGACAAGCTGGGCCGCAGCATTGTGCG
CAATTATCGCAAGAAGGTAAGAAGGCAATCAAGACTGACGGAAACCTCCGAACTATGCCTATCTATCATAT
ATATCCTGACAGCCTTCCCTGGTGGCCCGAGCGCCCAACACCAAGTATCCTTCGTTTCCGGAGTTTGTCA
ACTGGCTGCTCGACCAGGTGAAGCGGGGCAGCTTCATTGACATGCACTTTGTGGCGGCCACGTCGTTCT
GCACACCCTGTCTGATTCGTTTCGACATGATACTGAAGTTCGAGTCGCTCACAGAGGATCAATTATATCTA
ATCGAAAAGACTGGCCTGAAAAGGGTGATAGCGCCCGTGTGGCGCAACATGGGCAAAGGTGGCCGCAA
GACGCACGAACTCCAGCAGCAGTTCTATGCCCAGCTCACAAGGCACGAAATGCTGGAGCTCTACGAGTAT
TACAAGTGAGTGGGACAATCTAATACCCCTTTTGATAGTGCTCTAACAGTCTTTGTTCTATTCCAAGATATG

ATTTCGAGCTGTTTGACTACGATATCCAGGAGTATCTACAGGTAGCCAGACCAGACGAATCATCCGGCAG
TCCAACGGCCACAAAGAATTAACCCAGCTCTAAATTCGTAGAATTTGCTAACGAAATGAATTAGTTCAACG
AAAATACATATTAGCCGTCACAGTACGTAATTGCAAATGTTGATAATACTGAATGAATACAATAATAAACTTA
AATATATTTAAACGAAACGTTTTTACCGTCTATCGTTTTTTGGGCTGCTATCGATAAGCGGGATAACATTAT
ATCGAACAAATATCGGAACAAATATCGTTTGGGCCCATTTGGCATATTTGGAATTAGTGTGCCACAACAAC
TTTCTTCTGTTGCTAATTCCATGCACACATACCCTGGGGGAAATGAATGTTAAAAATTGAGAAAATGTTTGT
CTTCCAAGGCTCAGTAGGTATCAGGATCGCTATAAATATATACAAGATACTAATAATGTACTTGAATATGTC
TAAAGAATATCAATTTGAGAAAAGGTCTTTATAAATCACGTTTCATCTTCATATAAAATTAACAAAATAGGGT
ATACAAAGGCCAACACGCAACATAGAGACTCTTTTTTGTTGGACTTTTTCGTTTAAACCTTTTTTTACAATAA
ATACAATAAAAGCAAGGATTAGGTTAGGATAGGTTCAGGCGGTAGCCTGGGGGAGTGATGAAACCCTCCC
AAGCTCACTTGGACCTGTAGAAGGTCCGTTGTGGTGCCGCATGAGCGTGTGCCCCTTCCCAGTGCCTAAA
AACCGTGGAGAAACAGGCTGTTGCAAATTAAGGGCAGTCCCATCCGGAGGCTTGGATGAATCTGGACAA
GGTCCCCGGTTTGATTTCGGACAGGTCCGAAATGTCTGTGAGGAAAGCAGCCCCGAGAAAACAAAGACG
ACGATTTCCAAGGGCCGAGCAGTGTCAGAAGAAGTGGGGGACCGATTCCTCCTCTTCCTCGTCGCGACA
ACTCCTGCAGAAGTCGTTATGAGGGATTGACAGTCTAGAGGCATGAGTGCCAATCTGCCAGTGTCCCGTA
ATGGCACGAGTAACTGCAGAGCACTGTGCTCTGCTGAGCCTATACAGCTCGGCTGAGCGCTTCTTCGATC
GATATGGCCATATCAATCTTGAGATTTTACAGGAAATCTCCTATTGGCATTTTGCTCGAACAATTCGTGCGT
GAAGAGCCTGCACGTAGCCAGAGGCATCGCTACCTGCTCCCTCTCCATAAGGAGAGGAATTGTAGTACC
CTGCCTTGCTAGCTCGTCGGCGGCGTCGTTCCCCTCGATGTCCCTGTGGCCTGGGACCCATATAAGGAA
GAGGTCTAACTGATCTGCGATCTCGTGCAGAGACCTGCGGCAGTCATTTACAGTCGCTGAGTTCGACGAT
ATTGAGCCTAGAGCTTTAATAGCTACTTGGCTGTCCGAGAAAACGCACACTAAGTGGCGGTNGTCTNAGA
AGTAGTTGCAGACGATGGGAATGGGCCTCACTCCNACAGCGTCTGCGGGAATTAACCTGAATGCCTATAG
CAACATCGTCCGCATACGCCACCACACGGCAGCCACCCCCCTCTATCTCCCGCAGCAGTTCGTTCACTGC
CACGTTCCATAGGAGGGGCGAGAGGACTCCGCCCTGCGGGGTGCCTCTACTGACAAATCTGGTGGACGT
TGACGTCCCCAGTGATGCCTCAACCGTCCTGCATTGTAGCATCTGATCGATCAGTTTCACCGTCCGGGAG
TCAACCCCCAGGTCCGTCAGTGCACCCGTGATGGCGGTCGGGAGGATGTTATTAAAGGCGCCTTCTATG
TCGAGGAAGGCTACCAGGGTGTACTCCTTACAGTTGAGGGACGCTTCTATGATCGATGTGATCCTGTGTA
GGGCCGTTTCGGTGGATCTTCCTTTCCGATAGGCATGCTGGGAGTCTGAGAATAGACTAGGCGGAATATT
TACCGTGAGATGCATCCCCAAAAGCCGTTCCATCGTCTTCAGAAGGAAGGACGATAGACTTATGGGTCTG
AAATCTTTGGGGGCAGTGTGCGAGGGCTTGCCTGCCTTGGGGATGAAGACGACTTTGGGATGAAGACAG
GTTTCCGGGATATATCCATGGGAGAAGATTCCCTCGTATATCCTTTTGAGCCAATTAGTGGCTTNTTGTCC
AGCGTGAATCAGTTAGGGCAGGGATGATGCCATCTGGTCCCGCAGACTTGTATGGTTTGAAGCTTTTTATT
GCCCGGGCTCCTGGCTAAGCAGGTTCATGATAACACTTGAGGCAACGGAGGGAGCCGCGATGTAGTGTG
GTCTGTTTTCATCGCAGCCGGGAAAGTGGGTGTTAAGGAGAAGATTTAGCGACTCATTGCTGGACGTTGT
CCATGATTAGTCGGCGTTCTTCAGGTAGCCCAGAGTGGGTGTAGTCTTCGAGAGAACTTTGCGCAAGCGC
GAGGCTTCCGAGTTATTCTCAATGTCGCTACAGAACTTACGTCATGAGGCGCGTTTGGCTTTCCTCAGTTC
TTTATTGTAAACTGACAGACTGGCCATGTAATCGGCCCAGTTCTGCTCAGCGTTTTCAGCCTTATCTTTATT
GAAGAGTCTTCGGGAGGCTTTCCGGAGTTCCCTCAGTTTCGGATTCCACCATTCAGGTTTCTTCCGGCCT
CTGCTCTTGCTAGAGGGGCACGATCTATCGAGCGCGTCATTGCAGGCGTCGGTACATAGCTTAACAAGAT
GAGTCGTGGCTTCCCTGGAGATCTCCTCCTCAGGGAGAACACGACAGAGGTGGTCTGAGTACCGAGCCC
AGTTTGTCCGCCTAAGGTTCCGAAATGAGCTGGCCTCGGTAGTGAAAAAGAGAATACGGTTTCCACGTAC
CTGTGGTCCGAGAAGGAGTGCTCTTCCAGGACCCCCCAGGATACGATGTTCCCGTGTAACTCATGAGAG
GCAAGTGTGAGGTCCAGGACCTCCTTGCGGTTTTTAATAATGAAGGTGGGATCACTCCCCCTATTTAAAAG
GACCATCTGAGTGGTCAGTAAATAGTTGAAAAGGTACTCACCCCTTTCGTTGGTGTCAGTGCTTCCCCACT
GGCAGTGGTGTGCATTGGCGTCGCACCCTACGATTAGGCCGGTGTCCTTGGCCTCGCAGTCTATGATTA
GCTACGTATAGCTTATAGTCAGGAGTCCTCAGACCAGAGACCCTGTTTCCGAGGATCCAGGGCTCCTGAA
TGAGGACTATGTCGGCTCCACCCTTGGNCTAGNGTGGNGCAGGAGAGCAGCGCATGCTGCCTTACAAAG
GGTGGNGGTTTATCTGCAGGAGTATCAGTGACATTCCTGAGGTTCACCTCCACCATTGTCCTGTCGTGGT
CGCTTTCCGAAGAGTCAAACTCCTCCGCGACCCCGATGTGCTTGAGATCCCTAGTAAGATCGCTCACGTC
TGACACGTAACCATTAAGGATGTCATCCGACACCACTGATGCCACGTCTGACACCGCAGGCTTGATCTCC
ATACTAGGGATCTCCGGGAGCTCCAGGTCTGGCTCGACCGTCGGTTGCATGCCTTTGGAGTCGGACTTG

TACGGTACTACGACTACCTTCTTGTAGCCGTAGTCCCACAGCGCCCTCCGTGTCGTGGAGAAGTCTGACC
GACTCCTCGTTCAGGACGAGGATGATTTGGCGCCTCTGCCCGTTGCTCTCCTCTACCTTGGACACCTNCC
AATCGGCTGTGGGAAGGTGGGGGTTGCAGACCTGCAGAATCCGGAGAATCTTATCCGACTCTGCANNNC
TTCGCCGAGACCCACGCTCTGGCCCTCGGCTTGCTTCACTTCACGACCTCCAGCTGGGCTCCTGGGTAG
ACCTCCTTGAGCGACGCTACCGCCTTCGCGTAGAGGGCCGCCGAGCGAGCATGTTGGCAGGCGATGGC
TTTCACCCTGCCTTGGTGCCACCCGGCGTCCTCTCACTTGGGCGGTGGTCCTCCATTCTCCTCCAACTCC
TGGAGGAAACAGTCTTGGAGCTCGTTTTCCACGAGGTGCCACTTGTCCCTAGGGATCTGCCCGTCTTTAG
AGCCCCTGTCTAGGACTCCAATCAGGGTTTTTCCCCTCGCCACCTCGGCAAACGAGCGGTTGCTCAGTGT
CTTCGTCCTCTTGGCCTGTTGGACTTGTGTGGCGGTGTCCTCCGCCAGCTTTGGCTGGCTGGTAGTCGG
GCATGACTGTCTTGGCCCATCGCGGCGATTCGATTCCCTGGGCAGACGCCCGGAACACCGGGTCGTCGT
TGCCCAGGATGAAGGCCGCACGTCTTTTGTCCTGGAACTTGGGCCTATCTTTCGAGGCAGAGGCGCCGC
CTGCCGGGGTTGTCAAGGGGTTCCCGGTCCCTAGGGGTCCGCCAGCCGCGATATCGCTCTGCATGATCG
CCCCCCCGGTAACCGGGTCGCCCTTGGGGCCCCCGATGTGGATTGGCCCGATTGGCTTTTCGGTCCTCT
CCTCAAAGCGCCTGCTGCCTCGAGACGGTGGACCGACTTAGTCTCCTTCCCCGTCTTTTTGGGCACTGGT
TTCCAGATGCGTTGTAGAGGGATGGTCTTTCCTCCGGCACTTAGGAGGAGTGCCGTGCTCCTTGCGGTGT
TTTTAGTTTTAATATTAAATTTGGCATATTTGATCCCACGAGTAGGCGGGAAGGGGTTAGTCGCCCGAGCA
TAGCCCGCGTGCCCCGGGAAGCCTTATTAAAACTGGAGGTCGGCAGGTATCCAGAGTCCGCATACATAC
ATATGTATGTATGTATGTATGTATAAGCGACCGAGCACCCCCTCGGCCATGCATCCCTCGGCATATGAGA
GTGTCACCTTGGATTGGGGTTATTTCACTGAGAGTTTTCTTCTCTCCGCCCGACTACAATTAGCCAGTATC
CTGGCAGAGACATGACCGTACCAGGACCTGTTTCCAGCCGCCGTCACGGAGAGAGTATAGTCGCACTTC
CGCCGGTGTGTACAGTTACGGCGGGTGCGGGTTCGCTCGTGGCCACCTGTATCCTATGACGCGGAGCAC
AGTCGCCTTGGATCCAGGGCAAGCCATGAACCAGAGGCGCCTGTGCCCCGTTCCGAGTCTACAGTTGTG
ACGAGCCGACCCGACATCCGTTTATCCCCCTGTAGCACCCGACGGCTGACGGCCAAAAGCAAGGATTAA
AAACTAACTAATCTTGTAGAAAATTGACTCATCAATGGAATAAAACTATGTGGGCATTGCTGAGAGATTTAG
TTAGTCAGCATTATTCAATGGAATATCACAATTGAGCAATATGAACGAATTTCATTTTTCGTTTGTCTTGATT
GACCTATTTCGCTACTTGTTTTTTTGTTTGACTTATTTCGCTAAAACCCTTTTTCACGCAAAACGCAATAAAA
GCAAGTATTAAGAATACACCAGTTAAATAGAAAATTGATTCATCAATCGTATAAAATTATGTGGGCATGGTC
AAATTTTCTATCTAGCTAATAATATTCCTCGGAATATCAAAAACGAGGAATATGTAAAATAGAACTGGAATT
CGTCAGTATATTTAAGGTATATTTTAAAATGAGACGGTATATTTCTAAGGGTCGGACGGTATATTTTAACG
CGGTCAGACTGCAGCACTGCAACATCAACAAACGCCACGACCTATAAAAATTGTTAAATTAACTGACACCG
AAAATGAATGAGTAATTCAGTGAGAATCAATCGAAATTTATTAGCTAGAAGCATAGGCAAGGCATAAGCAA
AAATACTTACTTGTTTGTGATTTAGAGAGCTTCGGCATTGCACTGACGTGGACCTGAAAGATTAGAAACAG
GTGAGGCGACATGTGCATGCATTATCTAGATTGAAATCAGTCTGGTACACCTTTTGGCAGACTTCCAATAT
GTCCACAGCACGGCGGCGGCCCCAGGGTGGCAGTGCGGGCGATGTACAGAGCTACCACCCGAAGGAAA
GGCTGGTGGAGAAGGACGAGGGAACGCCGGACATACGCGGCGACAGACGGAACATAGCCATCCTGCTG
TTCCTGTATATACTGCAGGGCATACCCATCGGCCTGATAGCCGCCATTCCCATGCTGCTGCAGAACCGGG
GAGCCAGCTACAAGCAGCAGGCGGAGTTCTCCTTCGCCTACTGGCCGTTCAGCTTGAAGCTACTGTGGG
CCCCCATCGTGGACTCGCTTTACGTCCGGAGATTCGGGCGCCGCAAATCGTGGCTGGTGCCGGTGCAGT
ATCTGCTTGGCGGATTCATGATGTTCCTTTCCTACTACGTGGATCGTTGGCTGGGCGGCGATGGTGTGGA
GCCGAATGTGGCGCTGCTGTCGCTGCTCTTCTTCCTGCTCAATTTCCTGGCTGCCACCCAGGACATCGCT
GTGGACGCTGGCTCTGACTATGCTGAAGCGCTGCAATGTAGGCTACGCCTCCACCTGCAACAGTGTCGG
CCAGACAGCCGGCTACTTCCTTGGATATGTGTGTTCATTGCCCTCGAATCGAAGGACTTTTGCAACAAGTA
CATGAGGGATGTGCCCCTGAACGAGGGCATGATCACGTTACCACGCTTCCTCTGGTTCTGGGGTATTGTC
TTTGTGGTGGCCACCACTCTGGTGGCAATCTTCAAGAAGGAGAATGACATTGAAGATGCCCATACGGAAT
CTCGCTACACGGAGGAGCATGAGCTGAACATTCGTCAGAGCTACAAGATCCTCTGGGACATGGTGAAGAT
GCGACCAGTGCAGATCCTAGCCGCCATTCTGCTCACCGTTAAGGTAACCTTCTCCGCATCGGATGCGGTC
ACCAGTCTGAAGCTCATCGATGCCGGGGTGCCCAAGGATCAGCTGGCTCTGCTGGCCATCCCCCTCATT
CCCCTTCAGCTCGTCCTGCCCCTGGTGATGGGTCGCTACACCAACGGCCCGCGTCCCATGGATGTGTAT
CTGAAGGCCATTCCATATCGAATTCTTCTGGCCGCCGTGGCGACAATTTTCGCCTACGCCACACCTTTTAT
GGTGCAGAAAGGGCATGTCCCAGTGTACTATTATGTCCTTCTGATCGCCCTGTATGCCTGCTATCAGGTAT
TCCTGTACTCAATGTTCGTGGCGGCCATGGCATTCTTTGCAAAGATCTCGGACCCCGCTGTGGGTGGCAC

ATACATGACATTCCTCAACACGCTGTGTAATCTGGGCGGCAATTGGCCCAACACGGTGGTGCTCTGGCTG
GTGGACGTGCTCACATGGAAACAGTGCACCACCAATACGGATAATACGTGCCTCAATAAGGATGAGCAAC
AGGTACGAGAAGACTCTACCCACCTTCTGTGGTGTTTTTCAACTAATCTAGTGCCCTCCCTCACTCCTTAT
AGAGCTGTGAATCGTCACACGGCAATTGCGAGAT

### Supplementary Methods and Results

**Standard / Sex Ratio Females Fail to Produce Inversion Recombinants in cytogenetic mapping experiments.** Two isofemale lines one each collected from two localities (KBPN2, Kaibab National Forest, AZ, September 2017 latitude: 36° 24' 42" N; longitude: 112° 18' 48" W; AO4, Allred Orchard Farm Market, Provo, UT September 2017, latitude: 40° 15' 43" N; longitude: 111° 39' 32" W) produced all daughters suggesting each female had mated to an SR male. We crossed the putative ST/SR daughters to a hemizygous male with a multiply marked X chromosome. We individually crossed male offspring to ST/ST females (107 males from KBPN2 and 96 males from AO4). A single female larva (Figure S1) from each of the 107 KBPN2 and 96 AO4 male test crosses were karyotyped using standard cytogenetic methods (Painter, 1934). Specifically, we scored the XR chromosomal karyotype where all female offspring carried a ST chromosome derived from the F1 female parent plus the parental or recombinant chromosome from the tested male (Figure S2).

In the sample of 107 KBPN2 and 96 AO4 males from a ST/SR female, we found no recombinants among the three non-overlapping inversions. Thus, non-overlapping inversions are inherited as a single unit. The frequency of the ST gamete is not significantly different from the expected frequency of 50 % (see Table S3, KBPN2, binomial sign test $P$=0.281; AO4, binomial sign test $P$=0.063). The sex ratio for the ST arrangement was near 50% for both strains while SR males sired > 96 % daughters (Table S4). A t-test with unequal samples shows that the mean sex ratio is significantly different between ST and SR males (KBPN2, $t$=51.64, $df$=56, two tailed $P$ =$6.3 \times 10^{-49}$; AO4, t=33.73, df=91, $3.18 \times 10^{-53}$). Interesting, the mean number of offspring produced did not differ between ST and SR males (KBPN2, $t$=0.34, $df$=100, two tailed $P$ = 0.74; AO4, $t$=-0.00, $df$=92, two tailed $P$ = 0.99). Despite a reduction in Y-bearing sperm, SR males produce a similar number of offspring as a ST male.

**Table S1: *D. pseudoobscura* reference alignment statistics.**

| ***Statistic*** | ***D. pse ST*** | ***D. pse SR*** | ***D. mir*** |
|---|---|---|---|
| *Total Reads* | 139337556 | 145236658 | 49649299 |
| *Mapped Reads* | 133512488 | 139245407 | 45556023 |
| *% Mapped* | 95.82 | 95.87 | 91.53 |

### Table S2: *D. pseudoosbcura* reference alignment statistics for genomic scaffolds.

| Scaffold | Mean Coverage | Length | % Covered | Covered (bp) | + Reads | - Reads | Read GC | Median Coverage | St. Dev. Coverage |
|---|---|---|---|---|---|---|---|---|---|
| ***D. pseudoobscura ST*** | | | | | | | | | |
| 2 | 83.9129 | 30819483 | 97.5788 | 30073269 | 13841040 | 13841340 | 0.4403 | 85 | 40.51 |
| 3 | 85.9616 | 19787792 | 97.5288 | 19298803 | 9058245 | 9057166 | 0.4549 | 86 | 62.53 |
| XL_group3a | 44.7765 | 2692213 | 96.4761 | 2597342 | 642496 | 643846 | 0.4455 | 43 | 67.92 |
| XL_group3b | 54.3022 | 388551 | 98.341 | 382105 | 112865 | 113304 | 0.4266 | 45 | 122.17 |
| XL_group1e | 42.4095 | 12541198 | 97.8198 | 12267770 | 2878802 | 2874461 | 0.4426 | 42 | 26.38 |
| XL_group1a | 45.7357 | 9148293 | 96.8705 | 8861993 | 2257478 | 2255726 | 0.4523 | 42 | 81.27 |
| XR_group3a | 40.5871 | 1469181 | 97.7692 | 1436406 | 324078 | 324013 | 0.4676 | 40 | 16.46 |
| XR_group8 | 41.4123 | 9197557 | 97.3256 | 8951575 | 2056823 | 2057328 | 0.4571 | 41 | 20.03 |
| XR_group6 | 42.6727 | 13333775 | 97.5782 | 13010864 | 3061992 | 3063993 | 0.4477 | 42 | 34.97 |
| XR_group5 | 51.3656 | 740970 | 98.7634 | 731807 | 203154 | 203069 | 0.4408 | 44 | 49.9 |
| 4_group1 | 84.4553 | 5287126 | 98.1752 | 5190646 | 2402053 | 2400617 | 0.418 | 84 | 201.76 |
| 4_group2 | 81.5053 | 1235759 | 94.4519 | 1167198 | 534344 | 533721 | 0.4381 | 85 | 62.7 |
| 4_group5 | 80.7469 | 2439919 | 93.5238 | 2281905 | 1054607 | 1055036 | 0.4205 | 86 | 36.3 |
| 4_group3 | 83.7848 | 11685562 | 97.9635 | 11447582 | 5288757 | 5290633 | 0.438 | 84 | 38.18 |
| 4_group4 | 85.1283 | 6594820 | 96.8235 | 6385336 | 3016316 | 3013210 | 0.4307 | 86 | 58.84 |
| ***D. pseudoobscura SR*** | | | | | | | | | |
| 2 | 85.6435 | 30819483 | 97.5044 | 30050367 | 14142848 | 14139655 | 0.4399 | 87 | 39.7 |
| 3 | 87.4484 | 19787792 | 97.5056 | 19294208 | 9232111 | 9227540 | 0.4545 | 87 | 61.44 |
| XL_group3a | 46.4184 | 2692213 | 96.4888 | 2597685 | 666643 | 668042 | 0.445 | 45 | 70.15 |
| XL_group3b | 55.1312 | 388551 | 98.2872 | 381896 | 114618 | 114974 | 0.4253 | 46 | 102.94 |
| XL_group1e | 43.8398 | 12541198 | 97.8332 | 12269454 | 2978752 | 2973251 | 0.4421 | 43 | 27 |
| XL_group1a | 46.3822 | 9148293 | 96.8344 | 8858691 | 2295757 | 2290161 | 0.4523 | 43 | 92.78 |
| XR_group3a | 44.1206 | 1469181 | 96.2329 | 1413835 | 356858 | 355122 | 0.4634 | 42 | 60.14 |
| XR_group8 | 42.891 | 9197557 | 96.4071 | 8867101 | 2138713 | 2136598 | 0.4547 | 42 | 33.05 |
| XR_group6 | 44.1551 | 13333775 | 96.608 | 12881487 | 3185838 | 3187677 | 0.4467 | 43 | 50.19 |
| XR_group5 | 47.4404 | 740970 | 96.7479 | 716873 | 188068 | 188616 | 0.4374 | 45 | 39.7 |
| 4_group1 | 84.1611 | 5287126 | 97.9979 | 5181270 | 2393844 | 2394017 | 0.4193 | 87 | 60.4 |
| 4_group2 | 83.6336 | 1235759 | 94.4369 | 1167013 | 548437 | 547715 | 0.4372 | 87 | 58.68 |
| 4_group5 | 83.2656 | 2439919 | 93.3848 | 2278513 | 1087501 | 1088924 | 0.4196 | 88 | 44.79 |
| 4_group3 | 85.4123 | 11685562 | 97.8549 | 11434895 | 5396249 | 5400912 | 0.4376 | 87 | 40.35 |
| 4_group4 | 86.743 | 6594820 | 96.7652 | 6381492 | 3076471 | 3073661 | 0.4303 | 88 | 52.09 |
| ***D. miranda*** | | | | | | | | | |
| 2 | 18.7027 | 30819483 | 93.6237 | 28854340 | 4027647 | 4027315 | 0.4583 | 18 | 18.38 |
| 3 | 20.1168 | 19787792 | 94.4606 | 18691673 | 2779677 | 2785477 | 0.4691 | 19 | 26.53 |
| XL_group3a | 19.4302 | 2692213 | 92.6627 | 2494677 | 368119 | 368015 | 0.4623 | 18 | 32.81 |
| XL_group3b | 29.0281 | 388551 | 91.2452 | 354534 | 79627 | 79773 | 0.4578 | 19 | 47.92 |
| XL_group1e | 17.8448 | 12541198 | 92.6182 | 11615438 | 1585445 | 1585920 | 0.4625 | 17 | 21.8 |
| XL_group1a | 19.6939 | 9148293 | 91.9186 | 8408982 | 1275755 | 1276178 | 0.4701 | 18 | 44.68 |
| XR_group3a | 19.3349 | 1469181 | 93.2392 | 1369852 | 201744 | 200722 | 0.481 | 17 | 69.58 |
| XR_group8 | 17.3782 | 9197557 | 94.1218 | 8656902 | 1121618 | 1119342 | 0.4722 | 17 | 11.37 |
| XR_group6 | 18.1368 | 13333775 | 94.4924 | 12599410 | 1690148 | 1689856 | 0.4622 | 18 | 24.69 |
| XR_group5 | 20.3789 | 740970 | 94.8644 | 702917 | 105668 | 105158 | 0.4537 | 19 | 21.71 |
| 4_group1 | 16.3376 | 5287126 | 90.9867 | 4810580 | 609460 | 610129 | 0.4451 | 16 | 16.38 |
| 4_group2 | 18.8393 | 1235759 | 89.9682 | 1111790 | 162242 | 162151 | 0.4547 | 18 | 20.14 |
| 4_group5 | 16.2932 | 2439919 | 88.1169 | 2149982 | 279051 | 278681 | 0.444 | 17 | 14.46 |
| 4_group3 | 17.8253 | 11685562 | 93.7048 | 10949928 | 1461704 | 1460165 | 0.4556 | 17 | 23.03 |
| 4_group4 | 17.2507 | 6594820 | 89.696 | 5915288 | 802786 | 803197 | 0.4491 | 17 | 22.14 |

**Table S3. XR karyotype for tested male offspring from two ST/SR female strains.**

| XR Karyotype (female gamete/male gamete) | KBPN2 No. | Freq | AO4_No. | Freq |
|---|---|---|---|---|
| Par 1 $ST_1$ $ST_2$ $ST_3$/ $ST_1$ $ST_2$ $ST_3$ | 50 | 0.467 | 56 | 0.583 |
| Par 2 $ST_1$ $ST_2$ $ST_3$/ $SR_1$ $SR_2$ $SR_3$ | 57 | 0.533 | 40 | 0.417 |
| CO 1a $ST_1$ $ST_2$ $ST_3$/ $ST_1$ $SR_2$ $SR_3$ | 0 | 0 | 0 | 0 |
| CO 1b $ST_1$ $ST_2$ $ST_3$/ $SR_1$ $ST_2$ $ST_3$ | 0 | 0 | 0 | 0 |
| CO 2a $ST_1$ $ST_2$ $ST_3$/ $ST_1$ $ST_2$ $SR_3$ | 0 | 0 | 0 | 0 |
| CO 2b $ST_1$ $ST_2$ $ST_3$/ $SR_1$ $SR_2$ $ST_3$ | 0 | 0 | 0 | 0 |
| DCO 1 $ST_1$ $ST_2$ $ST_3$/ $ST_1$ $SR_2$ $ST_3$ | 0 | 0 | 0 | 0 |
| DCO 2 $ST_1$ $ST_2$ $ST_3$/ $SR_1$ $ST_2$ $SR_3$ | 0 | 0 | 0 | 0 |
| Total | 107 | | 96 | |

Par, parental; CO, cross over; DCO, double cross over;

**Table S4. Sex ratio (%female) variation across 107 and 96 F1 male offspring of a ST/SR female.**

| ST/SR Strain | ST | SR |
|---|---|---|
| KBPN2 | | |
| Range %female | 0.41 – 0.68 | 0.87 - 1.00 |
| Mean %female ± SD | 0.53 ± 0.06 | 0.99 ± 0.02 |
| Mean No. offspring ± SD | 121.9 ± 51.2 | 125.2 ± 47.1 |
| AO2 | | |
| Range % female | 0.47 – 0.84 | 0.80 - 1.00 |
| Mean % female ± SD | 0.56 ± 0.06 | 0.96 ± 0.05 |
| Mean No. offspring ± SD | 156.6 ± 39.9 | 156.6 ± 32.2 |

**Table S5: Counts from the recombination experiment. The reported recombination fraction is accompanied by exact 95% confidence intervals for the binomial distribution.**

| | Visible Marker Classes | | | | Recombination Fraction |
|---|---|---|---|---|---|
| Gene Arrangement | + + | + *sh*[1] | *se*[1] + | *se*[1] *sh*[1] | (95% Confidence Interval) |
| SR Chromosome Isolate 1 | 1459 | 0 | 3 | 1557 | 0.0010 (0.0002 - 0.0029) |
| SR Chromosome Isolate 2 | 1657 | 0 | 6 | 1914 | 0.0017 (0.0006 - 0.0036) |
| SR Chromosome Isolate 3 | 1464 | 0 | 3 | 1745 | 0.0009 (0.0002 - 0.0027) |
| ST Chromosome | 321 | 212 | 253 | 297 | 0.4294 (0.3996 - 0.4595) |

**Table S6: Primers used to amplify intergenic regions for linkage disequilibrium analysis.**

| | |
|---|---|
| **LD forward primer sequence** | |
| XL1_F | CTTTTGCGTGGGTGTGTTGC |
| XL2_F | TGCAACCGCACTTGACCGTA |
| XR1_F | ATGAGGGCGTTCCGAAAACAC |
| XR2_F | GTGTTTGGGTCGGGAACAGC |
| XR3_F | TGTCCCAGTCCCCGTTCTGT |
| XR4_F | AGCTGCCATCCCATTCCAAA |
| XR5_F | GGGCGAGACATGGGACATTC |
| XR6_F | TGCCTCGACCCACGAATACA |
| XR7_F | GCTGTTGCTGGGCAAACTGA |
| **LD reverse primer sequence** | |
| XL1_R | CGGGGACTCCTGCATTATCG |
| XL2_R | CTCGGCCAGAACCACATGCT |
| XR1_R | GCATTGGCCCCGAAAAATCAAC |
| XR2_R | AGCCGAACAGAACCGCAAAG |
| XR3_R | GCGGATTCGAACCATTCCTG |
| XR4_R | TAACTTGCACAGCCCCGTCA |
| XR5_R | CGGCTTCGGGAAACTTTGTGG |
| XR6_R | ACGGAACAAAACGGCCAAGA |
| XR7_R | AGCATCGCTTCGCATCTGTG |

**Table S7: RNA-seq read mapping data.**

| *Strain* | *Reads* | *Assigned* |
|---|---|---|
| KB10 | 96835955 | 83070610 |
| KB12 | 71249493 | 62675945 |
| KB1 | 48891591 | 42325411 |
| KB2 | 83033873 | 72656519 |
| KB3 | 33197393 | 29127193 |
| KB5 | 36001605 | 31805100 |
| NPZ11 | 93166191 | 80719581 |
| NPZ1 | 57669830 | 50083732 |
| NPZ29 | 68329286 | 58163461 |
| NPZ33 | 33991458 | 30042015 |
| NPZ6 | 95164635 | 82682359 |
| NPZ8 | 37619356 | 31948845 |

**Table S8: Reads per kilobase per million mapped reads (RPKM) and differential expression statistics for all genes.**

*See attached file DpseSR.GeneExpression.Data.xlsx*

**Figure S1**. *D. pseudoobscura* male and female larvae (Anterior to Posterior, left to right). The male larva has an obvious gonad about a third of the distance from the posterior end of the larva.

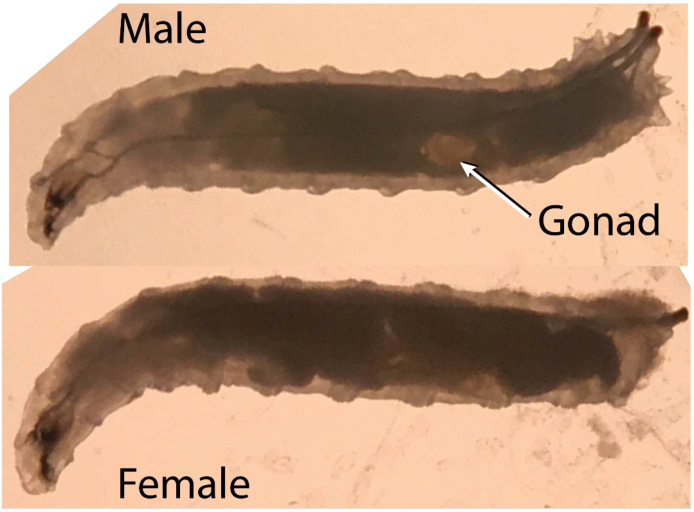

**Figure S2**. Genetic mapping crosses of KBPN2 and AO4 ST/SR females to detect cross overs among the three non-overlapping inversions that comprise the SR chromosome. Heterozygous ST/SR females were crossed to marked ST hemizygous males. F1 male offspring from each parental cross were individually crossed to virgin ST/ST females. Female larvae from each male were karyotyped at the three inversion loci. The F2 offspring were sexed and counted.

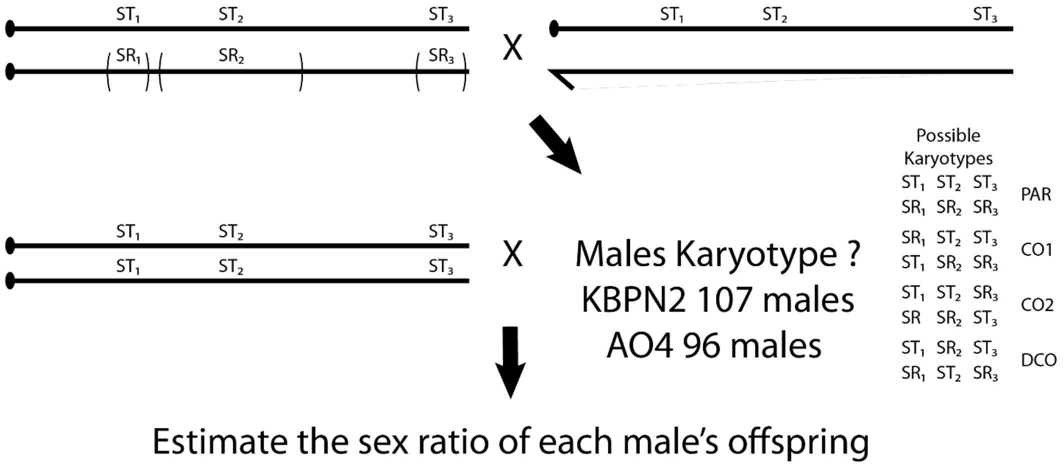

**Figure S3:** Structure of genetic variation on the autosomes in the RNA-seq data. A principal components analysis (PCA) was performed on genotypes called in all autosomal transcripts. The samples strongly cluster by ST/SR X-chromosome status, suggesting the presence of structured genetic variation present on the autosomes introduced by the crossing scheme used to generate the strains. As a result, only X-chromosome transcripts are analyzed further for differential expression.

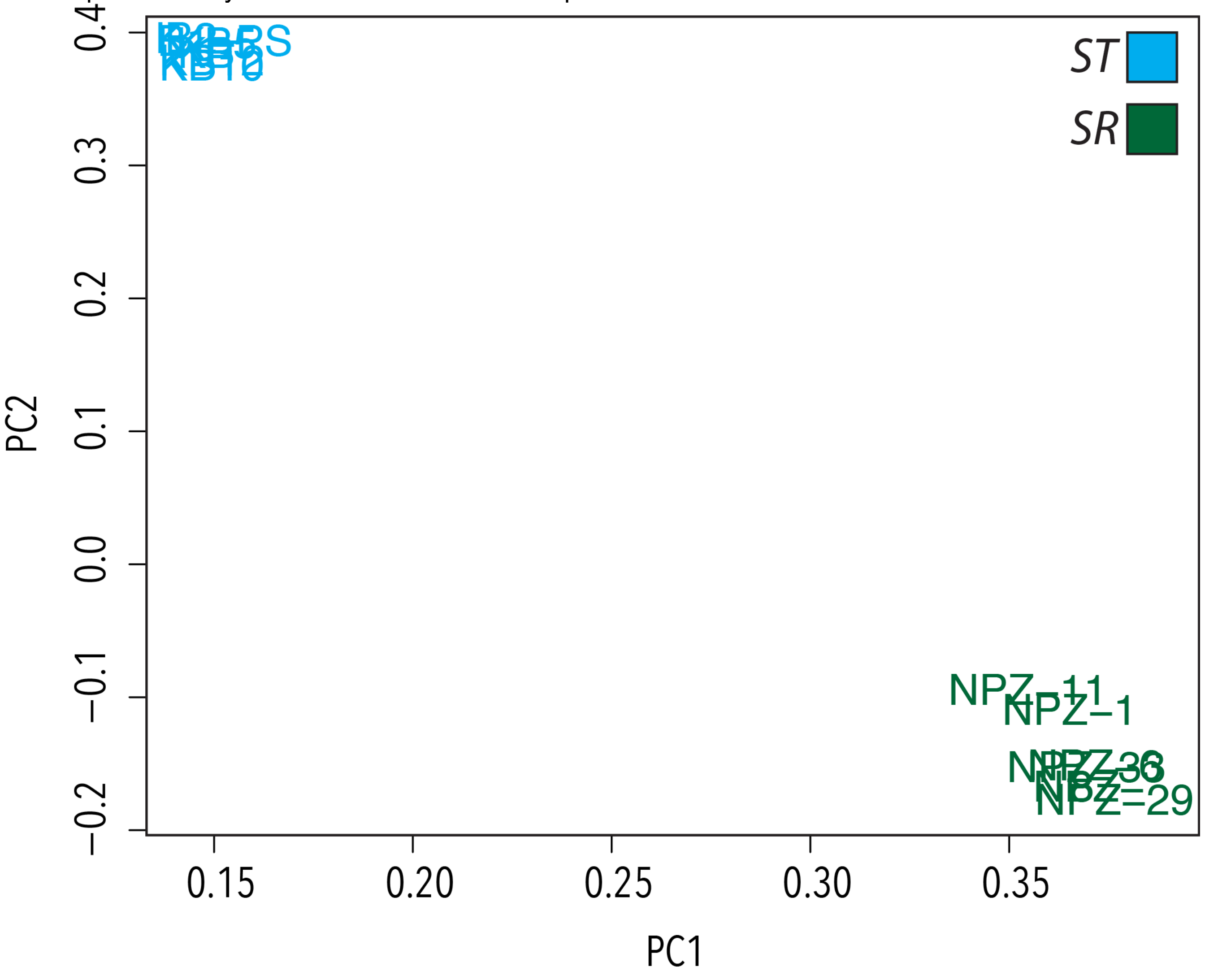
